# Accounting for biases in riboprofiling data indicates a major role for proline and not positive amino acids in stalling translation

**DOI:** 10.1101/006221

**Authors:** Carlo G. Artieri, Hunter B. Fraser

## Abstract

The recent advent of ribosome profiling – sequencing of short ribosome-bound fragments of mRNA – has offered an unprecedented opportunity to interrogate the sequence features responsible for modulating translational rates. Nevertheless, numerous analyses of the first riboprofiling dataset have produced equivocal and often incompatible results. Here we analyze three independent yeast riboprofiling data sets, including two with much higher coverage than previously available, and find that all three show substantial technical sequence biases that confound interpretations of ribosomal occupancy. After accounting for these biases, we find no effect of previously implicated factors on ribosomal pausing. Rather, we find that incorporation of proline, whose unique side-chain stalls peptide synthesis *in vitro*, also slows the ribosome *in vivo*. We also reanalyze a recent method that reported positively charged amino acids as the major determinant of ribosomal stalling and demonstrate that its assumptions lead to false signals of stalling in low-coverage data. Our results suggest that any analysis of riboprofiling data should account for sequencing biases and sparse coverage. To this end, we establish a robust methodology that enables analysis of ribosome profiling data without prior assumptions regarding which positions spanned by the ribosome cause stalling.

## Introduction

Translation of messenger RNAs into polypeptides by ribosomes is a fundamental process common to all life, and its dysregulation has been implicated in a number of diseases (Scheper et al. 2007). This has prompted a wealth of research into understanding the molecular underpinnings of translational dynamics. For instance, it has long been known that the frequency of codon usage in coding sequences (CDSs) is biased, suggesting the action of natural selection on the efficiency and/or accuracy of translational elongation (Kanaya et al. 2011; Plotkin and Kudla 2011).

The physiological origins of uneven codon usage have been studied extensively both experimentally and theoretically, implicating a number of different, non-mutually exclusive mechanisms - though all remain controversial (Plotkin and Kudla 2011; Gingold and Pilpel 2011). Much attention has been focused on the relationship between the cellular abundance of tRNAs and the frequencies of their cognate codons. Studies have found a strong correlation between gene expression levels and codon usage bias (CUB), revealing that highly expressed genes tend to use codons corresponding to the most abundant tRNAs in bacteria (Grantham et al. 1981), fungi (Bennetzen and Hall 1982), and metazoa (Shields et al. 1988; Stenico et al. 1994; Duret and Michiroud 1999) (though the abundances of charged tRNAs may be more important than total tRNA levels; Welch et al. 2009). As *in vitro* studies have shown that the rate of incorporation of amino acids varies in a codon-specific manner, with the most rapid rates of elongation occurring at codons with highly abundant tRNAs (Varenne et al. 1984), it has long been presumed that CUB reflects a general process of selection for high translational rate in highly expressed transcripts, preventing inefficient sequestration of ribosomes at slowly translated codons (Andersson and Kurland 1990). In addition, idiosyncratic requirements at individual mRNAs, such as induced ribosomal pausing to allow co-translational folding or shuttling, may also play a role (Thanaraj and Argos 1996; Corsi and Schekman 1996; Tsai et al. 2008).

Other factors thought to induce ribosome stalling include the presence of mRNA secondary structure, which must be ‘unwound’ by ribosomes, and may thus slow their rate of translation (Namy et al. 2006; Wen et al. 2008); wobble base-pairing, which can introduce non-optimal geometries in codon-anticodon interactions (Thomas et al. 1988; Kato et al. 1990); the presence of codons encoding positively charged amino acids, which may participate in electrostatic interactions with the negatively charged ribosomal exit tunnel (Lu et al. 2007; Lu and Deutsch 2008; Tuller et al. 2011; Charneski and Hurst 2013); and proline, which is inefficiently incorporated into polypeptides due to the unique structure of its imino side-chain (Wohlgemuth et al. 2008; Muto and Ito 2008; Pavlov et al. 2009; Johansson et al. 2011; Doerfel et al. 2013; Ude et al. 2013; Gutierrez et al. 2013; Zinshetyn and Gilbert 2013). Interpretation of the relative contributions of these factors has been challenging, as their effects have typically been studied in conditions not normally encountered in living cells – such as within genes with low CUB but extremely high mRNA levels (Plotkin and Kudla 2011; Gingold and Pilpel 2011).

However, this situation has changed radically with the recent development of ribosome profiling, an *in vivo* technique for monitoring transcriptome-wide rates of translation (Ingolia et al. 2009). By isolating and sequencing short fragments of mRNA bound by actively translating ribosomes, ‘riboprofiling’ provides nucleotide-resolution, quantitative information about the abundance and position of ribosomes on individual RNAs. When normalized for gene expression levels obtained by sequencing unprotected mRNA, increased ribosome-protected read coverage is expected from regions where ribosomes spend a greater fraction of their time, thereby identifying sequences that contribute to differences in rates of elongation (Ingolia et al. 2009; Ingolia et al. 2011).

Nevertheless, a number of recent studies that have analyzed yeast riboprofiling data have come to contradictory conclusions regarding the major determinants of translation rate, including whether non-preferred codons, RNA secondary structure, or particular amino acids stall translation (Tuller et al. 2010a, 2010b, 2011, Kertesz et al. 2010; Siwiak and Zelenkiewicz 2010; Zur and Tuller 2012; Qian et al. 2012; Charneski and Hurst 2013; Wallace et al. 2013; Rouskin et al. 2014). Unfortunately, direct comparison of these analyses has been challenging: while all have reanalyzed the *Saccharomyces cerevisiae* data of Ingolia et al. (2009), each study has made unique assumptions regarding interpretation of riboprofiling data – such as the precise location of active sites in ribosome protected fragments, or the effects of sequences near ribosome-protected fragments.

An additional consideration regarding the interpretation of riboprofiling data concerns the possibility that biases in the representation of particular nucleotides or sequences are introduced during library construction. For example, such biases have been well documented in the case of RNA-seq library preparation, where local base composition of RNAs can produce undesirable secondary structure, bias reverse transcription priming, and interfere with enzymatic steps such as ligation (Zheng et al. 2011). Such effects manifest themselves as protocol-specific biases in read coverage along transcripts, leading to over- or under-representation of certain sequences (Hansen et al. 2010; Srivastava and Chen 2010; Li et al. 2010; Bullard et al. 2010; Zheng et al. 2011). In studies of ribosome-protected fragments, such biases could confound identification of the actual biological factors affecting translational rate. However, the riboprofiling protocol itself provides a means to mitigate technical biases introduced during library construction: as the sequencing libraries generated from both unprotected mRNA (the ‘mRNA’ fraction) and ribosome protected mRNA fragments (the ‘Ribo’ fraction) differ only in the method used to isolate RNA, shared biases between the two are likely to represent technical artifacts (Qian et al. 2012).

In order to more thoroughly investigate factors that lead to increased ribosomal occupancy, we took advantage of two recently published yeast riboprofiling datasets that provide much higher coverage data than was previously available (Artieri and Fraser 2014; McManus et al. 2014) and compared them to the data of Ingolia et al. (2009). We observed consistent biased representation of specific nucleotides across all datasets that could be attributed to library construction. Controlling for these artifacts identified codons uniquely enriched in the Ribo fractions of the high-coverage datasets, suggesting that they may be responsible for ribosomal stalling *in vivo*.

## Results

### Riboprofiling data show consistent nucleotide biases

In order to explore how controlling for biases in library construction may affect our interpretation of sequence factors affecting translational rate, we analyzed two recently published *S. cerevisiae* riboprofiling datasets (Artieri and Fraser 2014; McManus et al. 2014); hereafter, the ‘Artieri and Fraser’ and ‘McManus et al.’ data (Supplemental Table S1). These data sets have ~28× and ~7× greater sequencing depth than was previously available (Ingolia et al. 2009), respectively. As most of the aforementioned analyses of ribosomal occupancy (Tuller et al. 2010a, 2010b, 2011, Kertesz et al. 2010; Siwiak and Zelenkiewicz 2010; Zur and Tuller 2012; Qian et al. 2012; Charneski and Hurst 2013; Wallace et al. 2013; Rouskin et al. 2014) used the *S. cerevisiae* data generated by Ingolia et al. (2009; ‘Ingolia et al.’ data), we also analyzed the raw reads from this study. The Ingolia et al. data include two different growth conditions: rich and amino acid starved media (hereafter ‘rich’ and ‘starved’; analysis of the starved data are in the Supplemental Material).

Reads from all samples were mapped to the *S. cerevisiae* genome (see Methods). Expression level estimates agreed well among replicates within each dataset (ρ = 0.96 - 0.99 and 0.92 – 0.99 for the Ribo and mRNA fractions, respectively) as well as between datasets (Spearman’s correlation coefficient ρ = 0.94 – 0.95 and 0.84 – 0.92 for the Ribo and mRNA fractions, respectively) (Supplemental Figs. S1 and S2). The Ribo fractions of all three datasets showed an enrichment of reads mapping at 28 – 29 nt, as expected based on the size of the ribosome-protected fragment (Ingolia et al. 2009); however the degree of enrichment varied among datasets (Supplemental Fig. S3; see Supplemental Material).

A larger proportion of Ribo fraction reads map to the first reading frame of codons as compared to the second or third (Ingolia et al. 2009), suggesting that there may be differences among reads beginning in different reading frames. Therefore we analyzed the nucleotide content of the first 27 nucleotides (nt) of reads, corresponding to the minimum mapping read length, from the mRNA and Ribo fractions separately for reads mapping to the first, second, or third frame of codons (Fig. 1). All datasets revealed substantial biases in the 5′ ends of reads that were similar in both fractions and among replicates, suggesting that certain sequences are preferentially incorporated during the process of library construction (Fig. 2, Supplemental Figs. S4 and S5). The most consistent of these biases is a preference for adenine in the 5′ most nucleotide, especially among first frame mappers. In the case of the Ribo fraction of the Ingolia et al. data, 66% of reads begin with adenine - two-fold greater than the adenine content within CDSs (32.6%). In comparison, 34% of the Artieri and Fraser and 33% of the McManus et al. Ribo fraction reads begin with adenine (Supplemental Table S2; see the Supplemental Material for a more thorough comparison of biases among datasets). The 3′ termini of reads in the Artieri and Fraser and Ingolia et al. datasets also showed a general preference for adenine, particularly in the mRNA fractions (Fig. 2). This is likely a consequence of the use of poly-adenylation as a template to prime reverse transcription; the McManus et al. data were generated with an alternative approach, which appears to mitigate 3′ adenine bias (see Supplemental Material).

**Figure 1.**
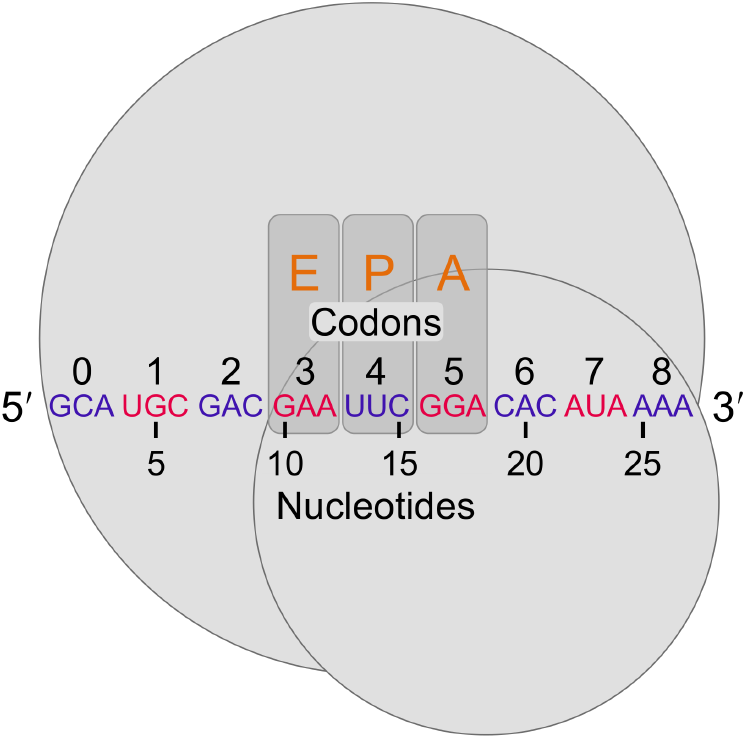
Defining positions relative to the 5′ end of riboprofiling reads. Following the mapping approach of Ingolia (2010), ribosomes (large and small subunits represented by grey circles) protect at least 27 nt of mRNA, corresponding to a minimum of nine codons. Both nucleotides and their overlapping in-frame codons were counted from 5′ to 3′ as shown (arbitrary codons are indicated in alternating blue and red for clarity). In the figure, the ribosome protected fragment begins in the first reading frame within a codon, and therefore codons are in-frame relative to nucleotides. However, for reads mapping to the second or third reading frames, while nucleotide counting begins at the first nucleotide, codon counting remains in-frame with the first codon, 0, corresponding to the one containing the first nucleotide. For reference, the orange letters indicate the codons that previous studies have indicated as the exit-tRNA (E-site), the peptidyl-tRNA (P-site), and aminoacyl-tRNA (A-site) sites, respectively (Ingolia et al. 2009; Stadler and Fire 2011; Qian et al. 2012; Li et al. 2012; Zinshteyn and Gilbert 2013).

**Figure 2.**
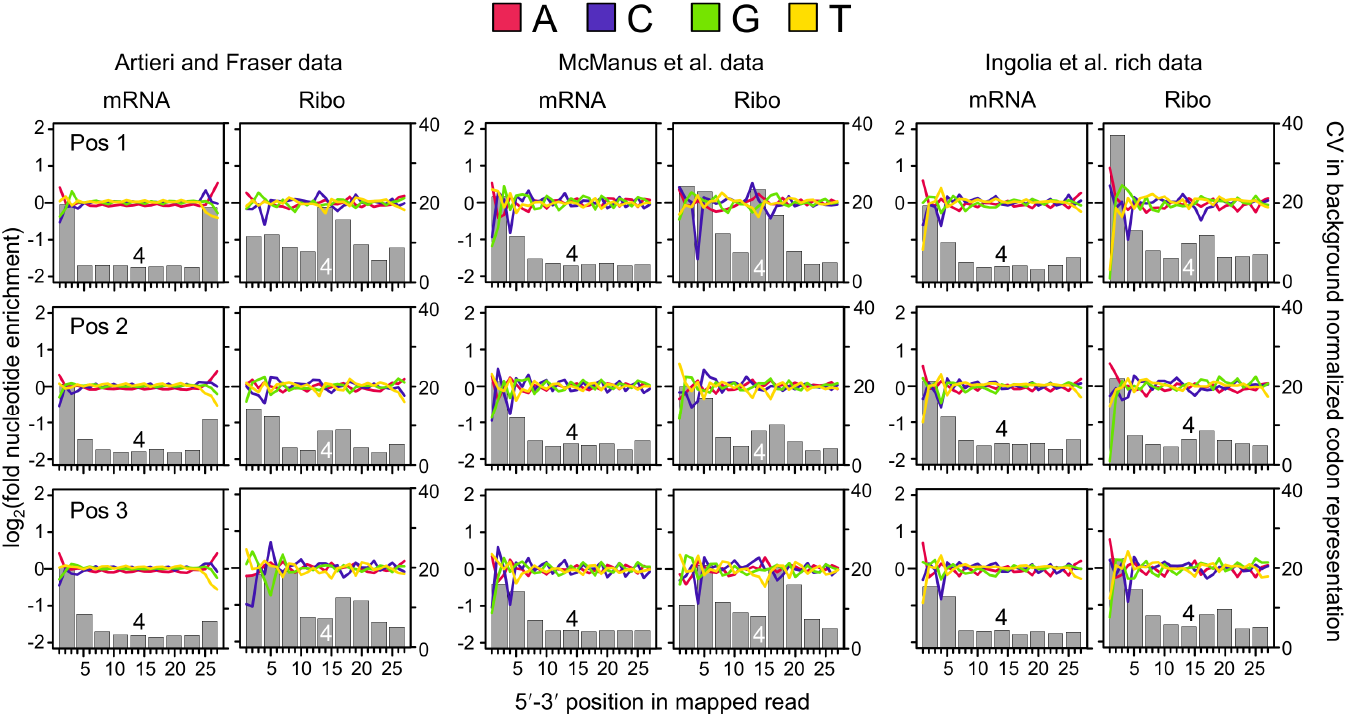
Patterns of nucleotide and codon representation across the three datasets. Reads were separated into those whose 5′ ends map to the first, second, or third reading frame within codons (frame 1, 2, or 3). The fold enrichment of each nucleotide was determined by dividing its number of counts at each position by the mean number of counts at positions within same reading frame across the 27 nucleotides analyzed, thereby accounting for differences in expected nucleotide proportions among reading frames within codons. Enrichment is plotted as colored lines in log_2_ scale: red, adenine; blue, cytosine; green, guanine; yellow, thymine. Each codon position overlapped by each read was also determined by identifying the nine consecutive codons beginning from the 5′ end as indicated in Fig. 1. The grey bars indicate the coefficient of variation (CV) as a measure of the degree to which each position deviates from the expected background frequency of the 61 sense codons; codon position 4 is indicated for reference. Both fractions in all datasets show similar degrees of nucleotide and codon biases at the 5′ ends of reads, though the biases in the Ingolia et al. Ribo data are generally more pronounced. The Ribo fractions show deviations from expected codons frequencies at internal read positions that are not shared by the mRNA fraction – in particular at codon 4 of first frame mappers. Third frame mappers in all datasets show a +1 codon shift in pattern. Note that the Ingolia et al. starved data show strong agreement with the rich data (Supplemental Fig. S4).

We assessed to what extent these nucleotide biases affected codon usage by identifying the nine consecutive in-frame codons spanned by each read (labeled positions 0-8 beginning from the 5′ end of mapping reads in Fig. 1) and determining the relative abundance of each of the 61 sense codons as compared to its expected frequency across all reads (see Methods). We then calculated the coefficient of variation (CV) among the relative abundances at each of the nine positions, producing a metric in which higher CVs indicate a greater deviation from expected codon frequencies (Fig. 2, Supplemental Figs. S4 and S5). Both fractions of all three datasets showed strong biases in position 0, as was expected from the observed nucleotide biases.

Interestingly, the Ribo fractions of the Artieri and Fraser and McManus et al. data showed strong biases at internal codon positions relative to the mRNA fraction – particularly in the case of first-position mapping reads – coinciding with the expected location of active ribosomal sites (Fig. 1), suggesting that these may reflect a biological signal of ribosome stalling. A similar pattern was observed in the Ingolia et al. data, though this was overshadowed by the stronger biases at 5′ codons (Fig. 2). We also noted that 28 nt reads, corresponding to the expected length of the ribosome protected footprint, showed stronger internal codon biases in all three datasets as compared to other mapping lengths (Supplemental Figs. S6 and S7; Supplemental Material). In contrast, the less common second frame mappers showed less pronounced internal codon biases. Interestingly, reads mapping to the third reading frame of codons in all three datasets were offset by +1 codon, indicating that the ribosome was likely positioned one codon downstream as compared to first and second frame mappers.

As first frame mappers are more abundant than reads mapping to the other two frames, and due to differences in 5′ bias and/or offset of reads mapping to the other two frames, we present subsequent analyses based on reads mapping to the first reading frame (however, similar results were observed for second- and third-frame mappers as well and are presented in Supplemental Material as indicated below).

### Ribosome occupancy is strongly associated with proline residues

Sequences that contribute to ribosome stalling should be enriched only in the Ribo fraction (Stadler and Fire 2011; Ingolia et al. 2011; Qian et al. 2012), whereas technical artifacts are likely to appear enriched in both fractions. Therefore, we normalized Ribo fraction coverage by that of the mRNA fraction (hereafter ‘corrected Ribo coverage’) as outlined in Fig. 3. Unlike previous studies (Ingolia et al. 2009; Stadler and Fire 2011; Qian et al. 2012; Li et al. 2012; Zinshteyn and Gilbert 2013), we did not attempt to define specific nucleotides corresponding to active ribosomal sites as we had no *a priori* expectations as to which position(s) near ribosome protected read fragments were responsible for stalling. Furthermore, we note that the hypothesis that positive amino acids stall ribosomes via interactions with the exit tunnel requires that they exert their effect after translation, and therefore should be enriched upstream of Ribo fragments (Charneski and Hurst 2013). Therefore we analyzed corrected Ribo coverage in a codon position-specific manner from eight codons upstream of the 5′ end of Ribo fraction reads to eight codons downstream (labeled positions −8 to +8, with position 0 corresponding to the in-frame codon to which the 5′ end of the read mapped). The log_2_-transformed enrichment of each codon at each of the 17 positions was scaled by the mean value of all codons at the same position such that codons with positive values were enriched relative to mean expectations and those with negative values were depleted (Fig. 3; see Methods).

**Figure 3.**
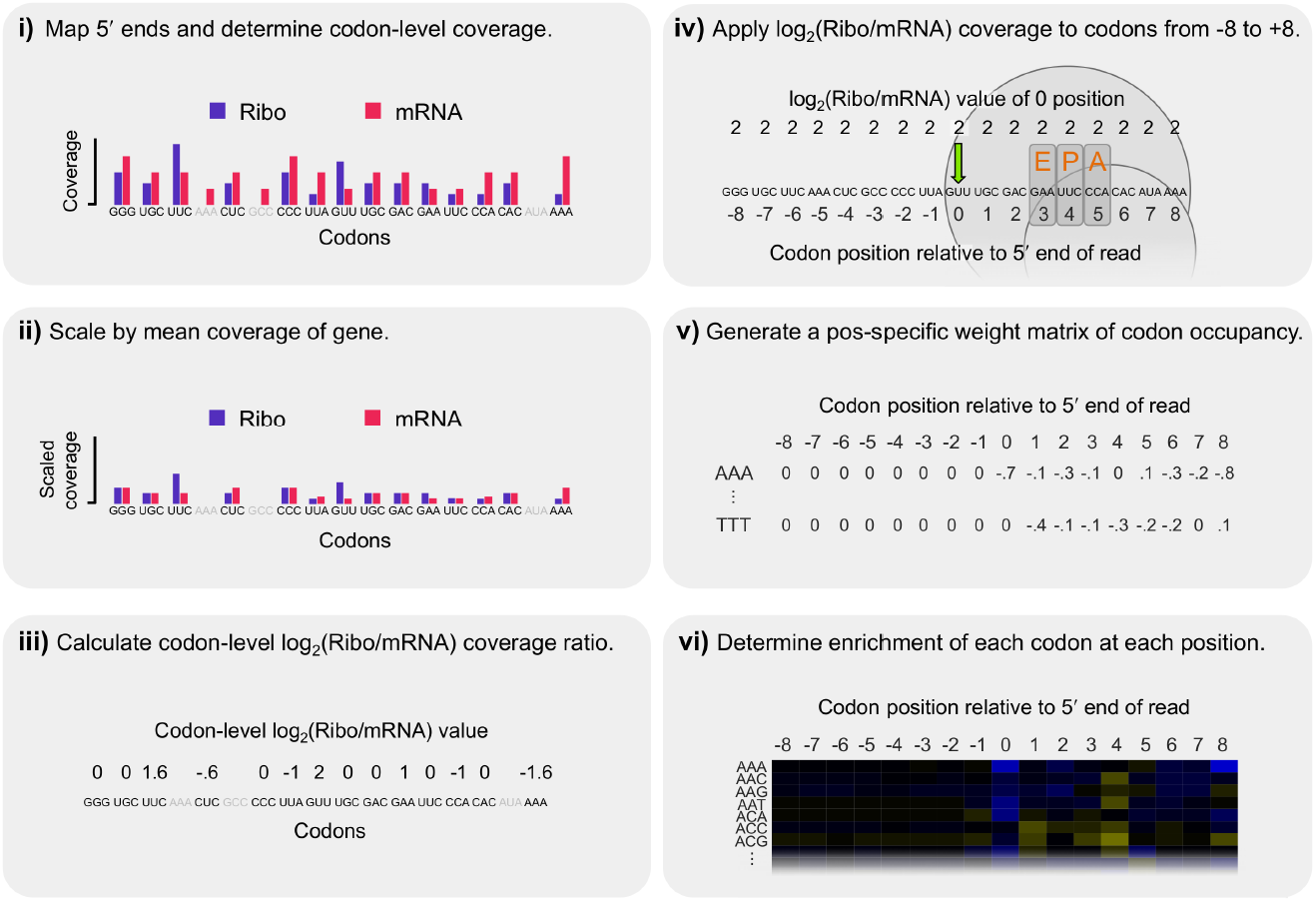
Steps in our calculation of corrected Ribo coverage. We analyzed Ribo fraction reads in a position-specific manner that controlled for shared biases between the two fractions while making no *a priori* assumptions about which codon position(s) may be most important in explaining patterns of coverage. **i)** The 5′ ends of reads were mapped and codon-level coverage determined from each fraction separately. Only sites with data from both fractions were considered (excluded codons are indicated in grey). **ii)** To account for coverage differences among genes, codon-level coverage values were scaled by the mean codon-level coverage of analyzed sites within each gene. **iii)** These scaled values were used to calculate a log_2_(Ribo/mRNA) coverage ratio for each codon, thereby accounting for shared biases between the two fractions. **iv)** As increased coverage at the 5′ position of ribosome protected fragments could be driven by sequence factors up- or downstream, the log_2_(Ribo/mRNA) coverage was applied to all codons from −8 to +8 codons relative to the 5′ end for each analyzed site (position ‘0’ shown by the green arrow). The expected position of the ribosome is indicated for reference. **v)** In this manner, the mean log_2_(Ribo/mRNA) value for each codon at each position was determined, generating a position-specific weight matrix representing each codon’s average occupancy at each of the 17 positions. **vi)** Finally, the relative enrichment of each codon at each position was determined by scaling its mean log_2_(Ribo/mRNA) coverage value by the mean value of all 61 sense codons at that position such that codons with positive log_2_ values were enriched relative to expectations and those with negative values were depleted (as plotted as in Fig. 4A).

The scaled enrichment values of all 61 sense codons at positions −8 through 8 as determined from the Artieri and Fraser data are shown in Fig. 4A, revealing that the strongest enrichment occurred at position 4, which corresponds to the position of the elevated internal biased codon representation observed in Ribo fraction reads (Fig. 2). In addition, as compared to positions spanned by the ribosome-protected fragment (i.e., 0 to 8), there is no evidence of substantial enrichment of upstream codons (−8 to −1) as would be expected if positive amino acids slow translation as they pass through the negatively charged ribosome exit tunnel (Lu et al. 2007, Charneski and Hurst 2013). Near identical results were also observed in the McManus et al. data (Supplemental Fig. S8). Therefore the strongest single-codon stalling effect was driven by codons in the position defined by previous studies as the P-site and not the A-site (position 5) (Ingolia et al. 2009; Stadler and Fire 2011; Qian et al. 2012; Li et al. 2012; Zinshteyn and Gilbert 2013). Reads that mapped to second and third codon frames showed qualitatively consistent patterns at upstream and downstream codon positions, though the degree of enrichment varied (Supplemental Fig. S8). This pattern disappeared completely in both datasets when the order of codons was randomly shuffled within each gene, preserving the relative position of mapped reads, indicating that it was not an artifact of the relationship between codon order and patterns of read mapping positions within transcripts (Supplemental Fig. S9).

**Figure 4.**
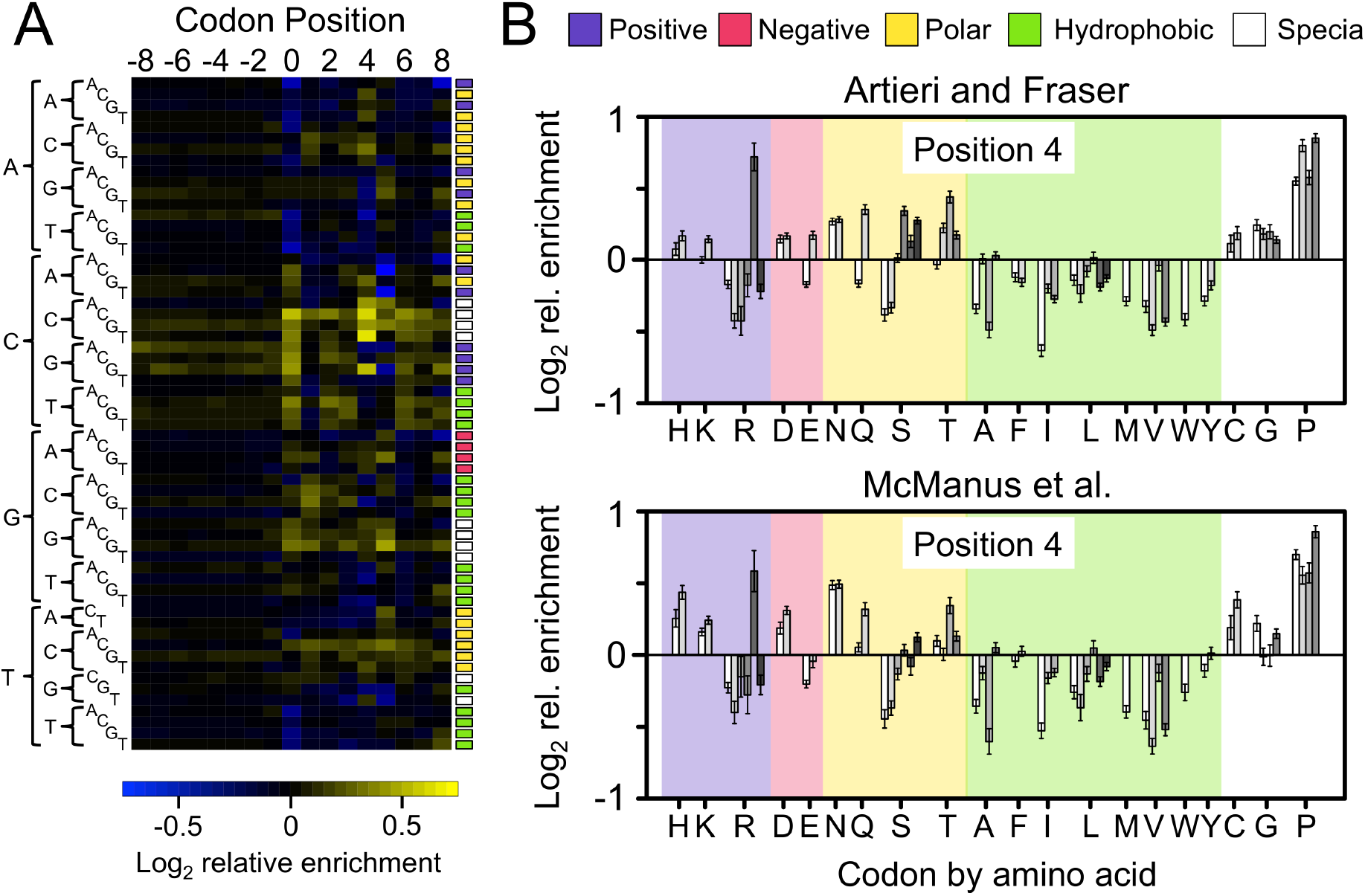
The corrected Ribo coverage reveals a strong enrichment of proline codons. **A)** Heatmap of the mean-scaled log_2_ enrichment of codon positions −8 to 8 in the Artieri and Fraser data (the McManus et al. data are qualitatively similar; Supplemental Fig. S8). All 61 sense codons are shown in alphabetical order indicated by their sequences on the left. Enriched codons are indicated by an increasing intensity of yellow color while depleted codons are blue. For reference, colored boxed to the right of each row indicate the biochemical category to which the codon belongs (color-category correspondences are indicated at the top of panel **B)**. Codon position 4 is an outlier in terms of enrichment. **B)** Bar plots indicating the log_2_ enrichment values at position 4 of both the Artieri and Fraser and McManus et al. datasets. Codons are organized by amino acid using single-letter designations below and grouped by biochemical type as indicated at the top of the panel. Individual codons for each amino acid are in alphabetical order. 95% confidence intervals around the scaled enrichment values are indicated at the top of each bar. No significant enrichment is observed among all codons belonging to a particular biochemical category, while a general paucity of representation is observed for the hydrophobic category. However, proline (P) codons are among the most highly enriched in both datasets, and are significantly more enriched than any other amino acid (Kruskal-Wallis rank sum test, p < 10^−15^).

Unlike the other datasets, the magnitude of 5′ bias observed in the Ribo fraction of the Ingolia et al. data was significantly stronger than that observed in the mRNA fraction (see above; Fig. 2), and consequently a much stronger 5′ codon bias remained after normalization by the mRNA fraction, overwhelming any patterns observed at internal codons (Supplemental Figs. S10 and S11). Therefore, we restricted our analysis of factors affecting ribosomal occupancy to the two higher-coverage datasets and discuss the two Ingolia et al. datasets in the Supplemental Material.

As position 4 showed the strongest degree of preference for particular codons among internal positions (Figs. 2, 4A), we analyzed patterns of codon enrichment at this position. We first explored whether any biochemical properties of amino acids (i.e., positive, negative, polar, or hydrophobic) were significantly enriched (Fig. 4B). No category showed consistent enrichment, including positively charged amino acids, which were recently implicated in stalling based on a reanalysis of the Ingolia et al. riboprofiling data (Charneski and Hurst 2013; see below). However, both datasets did show a general paucity of coverage among codons for hydrophobic amino acids.

Among individual amino acids, both datasets showed a higher level of enrichment among proline codons (CCN) than for any other amino acid (Kruskal-Wallis rank sum test, p < 10^−15^) (Fig. 4B). The four proline-encoding codons were among the five most enriched codons in both datasets (the fifth, CGG, encodes arginine; see below). These results were reproducible among different subsets of mRNA expression levels, indicating that they were not driven by highly abundant genes (Supplemental Fig. S12). Furthermore, they are consistent with proline’s previously implicated role in translational pausing *in vitro* (Wohlgemuth et al. 2008; Pavlov et al. 2009; Johansson et al. 2011).

We also tested whether other specific factors were associated with increased ribosomal occupancy. Previous riboprofiling studies have suggested that mRNA secondary structure can slow translation (Tuller et al. 2010b, 2011). Therefore, we looked for evidence of increased corrected Ribo coverage upstream of regions of mRNA secondary structure (Ouyang et al. 2013). However, we observed that terminal adenine biases within both riboprofiling datasets had stronger correlations with measurements of secondary structure than any potential signal of ribosome stalling (Supplemental Fig. S13; Supplemental Material). In addition, G:U wobble base-pairing has been associated with pausing in nematodes and humans (Stadler and Fire 2011). Although we observed no such pattern at codon position 4, Watson-Crick pairing was enriched at position 5 (Supplemental Fig. S14). Nevertheless, the magnitude of this enrichment was relatively modest, indicating that it is not a major determinant of ribosomal stalling in yeast.

Finally, supporting previous riboprofiling-based observations made in yeast Qian et al. 2012; Zinshteyn and Gilbert 2013), *E. coli* (Li et al. 2012), and mouse (Ingolia et al. 2011), we found no correlation between corrected Ribo coverage and non-optimality of codons at either position 4 (P-site) or at position 5 (A-site) using three separate measures of codon optimality (Supplemental Figs. S14 and S15; Supplemental Material). Interestingly, the rarest codon in *S. cerevisiae*, CGG (encoding arginine), showed a substantial level of enrichment in both datasets (Fig. 4B). However, this may not be related to its rarity, as similarly rare codons (CGC and CGA, also encoding arginine), showed no such enrichment.

### Revisiting the effects of positively charged amino acids

A recent reanalysis of the Ingolia et al. data concluded that positively charged amino acids were the primary determinant of ribosomal velocity (Charneski and Hurst 2013). Their approach assumed that upon encountering a sequence feature causing ribosomal stalling (such as a positive amino acid), the ribosome slows, leading to an accumulation of Ribo fraction reads immediately downstream of the feature. By comparing the magnitude of this accumulation to read coverage upstream of the stalling sequence – where the rate of translation was presumed to be unhindered – they generated a normalized metric of stalling as shown in Fig. 5. Specifically, to test the effect of a codon at position 0, the occupancy of all codon positions (r_pos_) from 30 codons upstream to 30 codons downstream was divided by the mean occupancy of upstream codons −30 to −1 (r_prec30_), producing a normalized pausing value (r_pos_/r_prec30_) where a value of 1 represents the average rate of translation. The area under the curve (AUC) of the mean-normalized occupancy values from position 0 until the position where mean occupancy returned to the average was used as a measure of the stalling effect, if positive (Fig. 5).

**Figure 5.**
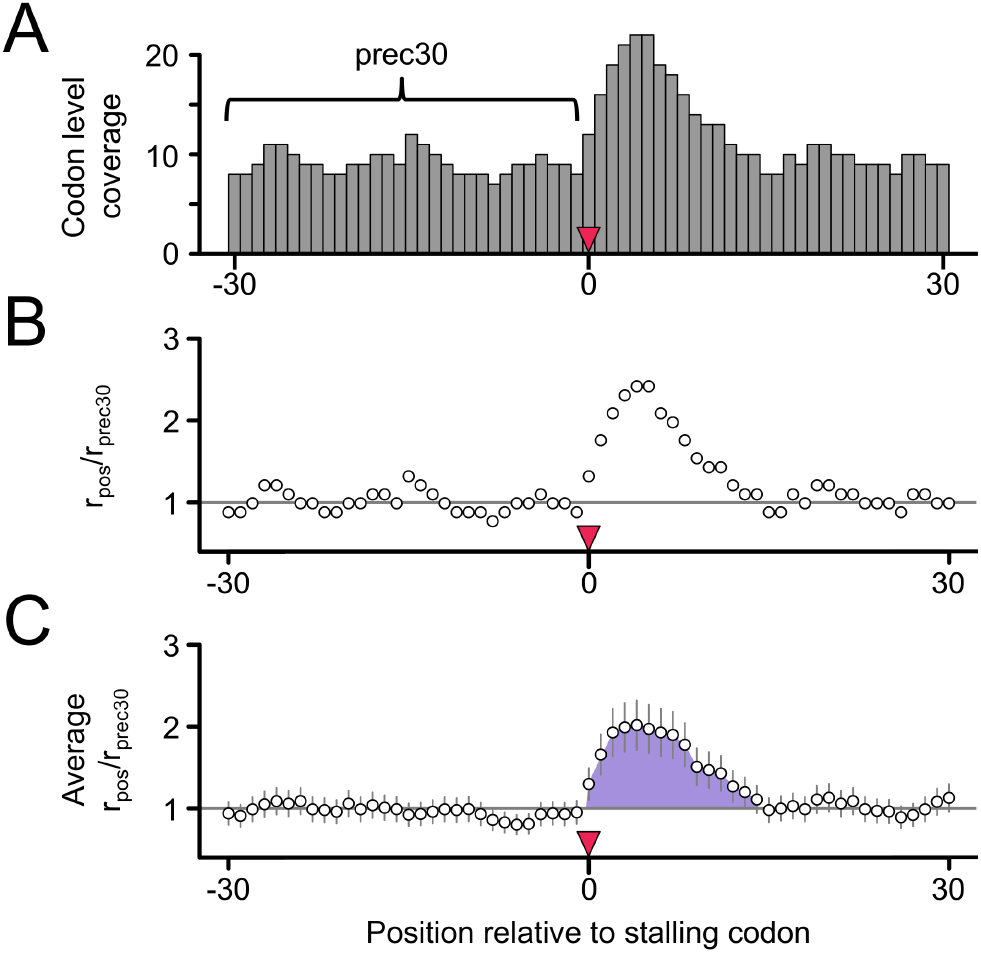
The r_pos_/r_prec30_ method of Charneski and Hurst. As a measure of the stalling effect of a codon (or group of codons beginning) at position 0, **A)** the occupancy of all codon positions (r_pos_) from 30 codons upstream (position −30) to 30 codons downstream (position 30) of the putative stalling codon was divided by the mean occupancy of upstream codons −30 to −1 (r_prec30_, indicated by the bracket). **B)** This produced a normalized pausing value (r_pos_/r_prec30_), where a value of 1 represents the average rate of translation. **C)** After averaging the r_pos_/r_prec30_ values among all similar groups of codons, the AUC (indicated by the shaded blue area) of the mean-normalized occupancy values from position 0 until the position where mean occupancy returned to the average was used as a measure of the stalling effect (if positive).

We sought to test if the stalling effect of positive amino acids was also detected in the higher coverage Artieri and Fraser and McManus et al. datasets. We first replicated the additive pattern of increased stalling with increasingly large clusters of positive amino acids (Figure 5 in Charneski and Hurst 2013) using the Ingolia et al data, confirming that the same methods were being used (Supplemental Fig. S17). However, analysis of both higher-coverage datasets showed no such coherent additive stalling trend (Fig. 6A-B; for one, two, three, four or five, and six or more positive charge clusters, the AUCs for the Artieri and Fraser data were 7.89, 12.83, −0.71, −1.36, and −2.75, and for the McManus et al. data were 6.46, 0.08, −0.59, 0.04, and 0.09, respectively).

**Figure 6.**
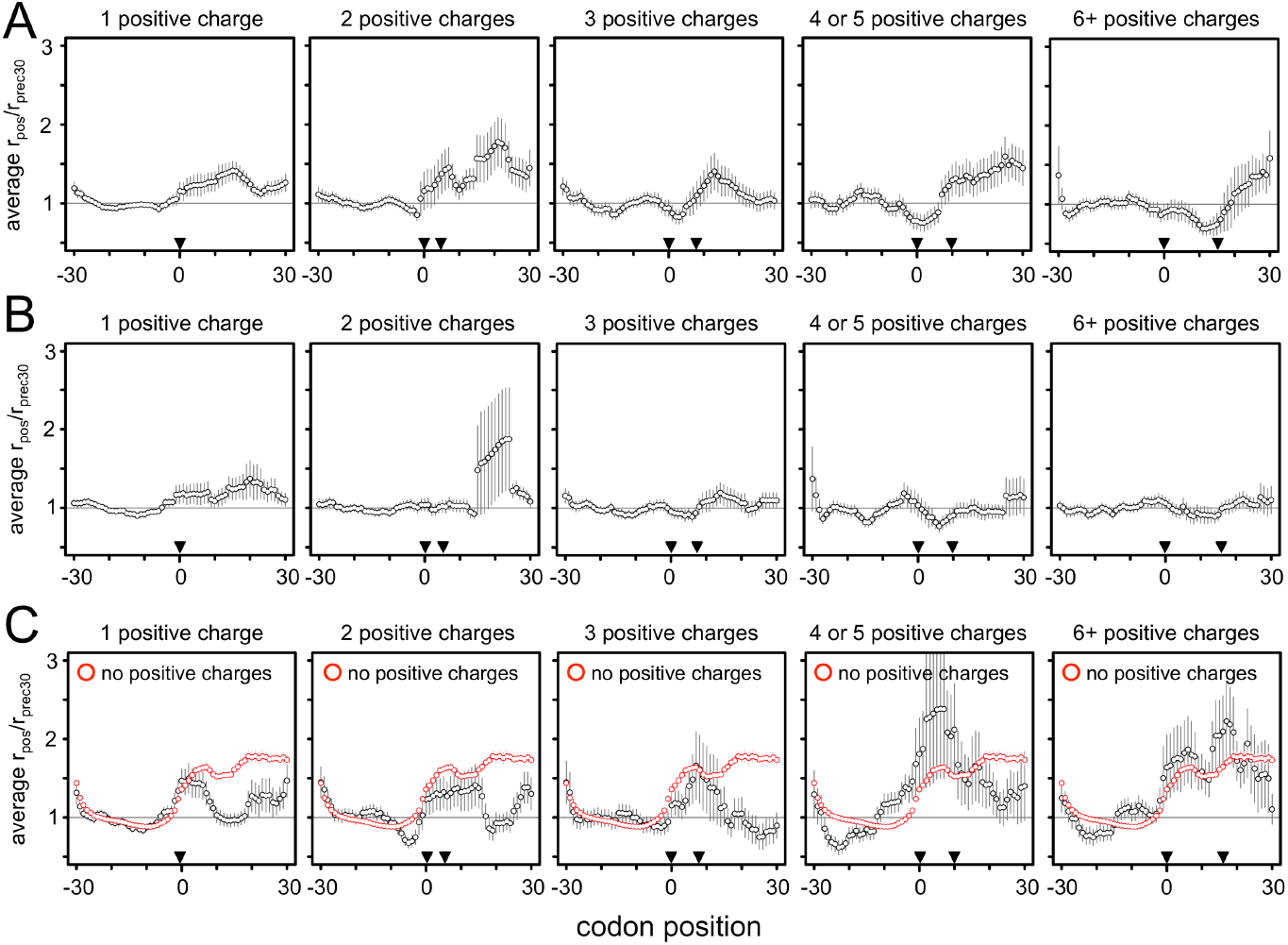
No evidence of stalling at positive amino acids. We recalculated Charneski and Hurst’s (2013) Figure 5 using either **A)** the Artieri and Fraser or **B)** the McManus et al. data. All analysis of mapped reads was performed as in the original manuscript, where clusters of increasing numbers of positive amino acid encoding codons were identified within the range bounded by pairs of inverted triangles. The horizontal gray line indicates the average rate of translation. The error bars represent ± the standard error of the mean. No additive effect is observed in either high-coverage data set, in contrast to the Ingolia et al. data (Supplemental Fig. S17). **C)** The data from Charneski and Hurst (2013) Figure 5 (black) compared to the mean r_pos_/r_prec30_ generated from 100 random samplings of 61-codon windows devoid of any positive amino acid encoding codons (red). The average stalling pattern of windows lacking any positive charges is stronger than clusters of one to three positive charges and is not significantly different from clusters of more than three. Therefore the observed stalling effect of positive amino acids is not greater than what would be expected by chance within the Ingolia et al. data.

Analysis of coverage of either the corrected or uncorrected Ribo coverage showed no systematic pattern of enrichment among positive amino acids among upstream codons (position −8 to −1) in any of the three datasets (Supplemental Figs S8, S10, and S18; Supplemental Material). Therefore, we explored the r_pos_/r_prec30_ method in more detail by performing an important control not reported in the original analysis (Charneski and Hurst 2013): levels of apparent stalling in the absence of any positive amino acids, using the same data set (Ingolia et al. 2009) (see Materials and Methods). We found that the median apparent stalling effect was actually *stronger* in the absence of any positively charged residues than in any sized clusters of positive charges (Kruskal-Wallis rank sum test of distributions AUC values, p < 10^−15^ for all clusters except for 6 or more positive charges, where p = 0.02 after Bonferroni correction for multiple tests) (Fig. 6C). We observed a similar pattern of stalling when averaging over all possible 61-codon windows in all genes (Supplemental Fig. S19), suggesting that the apparent pattern of stalling is unlikely to be related to the presence of positively charged amino acids.

We then explored whether read coverage could affect these patterns even in the absence of any stalling by generating simulated data at a range of coverage levels. Indeed, stalling was observed in low-but not high-coverage windows (Supplemental Fig. S19; Supplemental Material). Since the simulated data contained no actual stalling, we concluded that the r_pos_/r_prec30_ method will detect stalling in any series of windows with sparse read coverage. As a further test, we downsampled the higher-coverage data to the level used in the original analysis, and found that overall patterns of stalling indeed increased (Supplemental Fig. S20).

## Discussion

### Library construction biases

The relative importance of various factors implicated in influencing the rate of translation has remained controversial despite recent advances in our ability to measure translation rates at the level of individual codons (Plotkin and Kudla 2011; Gingold and Pilpel 2011). Most of these factors were originally identified using *in vitro* approaches, which may not accurately represent intracellular conditions. As an *in vivo* method, riboprofiling has offered an unprecedented opportunity to study translational dynamics in living cells; yet a number of different studies reanalyzing the same riboprofiling data (Ingolia et al. 2009) have produced incompatible findings, based on differing assumptions and methods of analysis (Tuller et al. 2010a, 2010b, 2011, Kertesz et al. 2010; Siwiak and Zelenkiewicz 2010; Zur and Tuller 2012; Qian et al. 2012; Charneski and Hurst 2013; Wallace et al. 2013; Rouskin et al. 2014).

Our approach presents a number of improvements over previous analyses of the biological basis of increased ribosomal occupancy using riboprofiling data: First, we have explicitly taken into account shared technical biases between the Ribo and mRNA fractions. Second, we made no *a priori* assumptions regarding which codon positions near the ribosome-protected fragments were responsible for rate variation, but rather focused on codon position 4 because it was a clear outlier in terms of enrichment in corrected Ribo coverage. And third, we analyzed two independently generated, high-coverage datasets (Artieri and Fraser 2014; McManus et al. 2014) and found strong agreement between them.

Our analysis revealed that like other next-generation sequencing methods (Hansen et al. 2010; Srivastava and Chen 2010; Li et al. 2010; Bullard et al. 2010; Zheng et al. 2011), riboprofiling is subject to library construction biases that may confound any analysis of ribosomal occupancy. In particular, both fractions of all three datasets showed a substantial preference for adenine bases at the 5′ ends of reads (and in some instances, the 3′ ends as well) (Fig. 2; Supplemental Material). As the majority of reads from the Ribo fraction mapped to the first reading frame of codons, this produces skewed representation of reads mapping to codons that begin with these bases. In the Ingolia et al. data in particular, the biases at the 5′ ends of reads overwhelmed those of all other positions spanned by the reads, suggesting that patterns of Ribo fraction coverage are strongly influenced by this library construction bias (Fig. 2; Supplemental Fig. S18; Supplemental Material). Also, we note that an additional caveat applicable to all yeast riboprofiling datasets discussed in this manuscript is that cycloheximide was used to arrest translation immediately prior to RNA extraction (Ingolia et al. 2009; Zinshteyn and Gilbert 2013; Artieri and Fraser 2014; McManus et al. 2014). It is unknown whether the drug itself has any sequence-specificity, but if so, this could lead to artifactual signals of ribosome stalling.

### Proline codons are enriched in corrected Ribo coverage

Of the features previously implicated in modulating the rate of translation, we observed consistent enrichment of Ribo coverage only at proline residues (Fig. 4B): all four proline codons (CCN) were among the most significantly enriched at codon position 4 in both the Artieri and Fraser and McManus et al. data. Interestingly, position 4 corresponds to what previous studies have defined as the P-site (Ingolia et al. 2009; Zinshteyn and Gilbert 2013; Stadler and Fire 2011; Li et al. 2012), where the imino side-chain of proline is known to act as a particularly poor substrate in the peptidyl transfer reaction. This is likely due to its restricted conformational flexibility, which may limit the rate of translational elongation (Wohlgemuth et al. 2009; Pavlov et al. 2009). Proline’s ribosomal pausing effect is known to play an important role in programmed stalling (Gárza-Sanchez et al. 2006; Tanner et al. 2009), and previous riboprofiling studies have found an enrichment of proline codons in the context of multi-amino acid motifs (PPE, Ingolia et al. 2011, and PG, Zinshteyn and Gilbert 2013), even in the context of cells untreated with cycloheximide (Ingolia et al. 2011).

Several other amino acids showed a lower enrichment among their corresponding codons in both datasets (Fig. 4B); however, these were each encoded by just two codons, making it difficult to determine if this is due to properties of the amino acid residues or the codons themselves.

### No evidence that positive amino acids stall ribosomes

Though several recent studies have suggested that positively charged amino acids may impede the progress of the peptide chain through the negatively charged ribosomal exit tunnel (Lu et al. 2007; Lu and Deutsch 2008), we observed no consistent enrichment for codons encoding positive amino acids in corrected Ribo coverage either within or upstream of the footprints in any datasets (Fig. 4B; Supplemental Fig. S18). Two previous studies found an association between riboprofiling read coverage and the presence of positive amino acids in yeast – both based on reanalysis of the data of Ingolia et al. (2009). The first (Tuller et al. 2011) noted an association between increased ribosomal occupancy at the 5′ ends of CDSs and an increased incidence of positive amino acid encoding codons. However this increased frequency of positive amino acids can be explained entirely by the requirements of hydrophilic N-termini of transmembrane proteins (Charneski and Hurst 2014). With regard to the second study, that of Charneski and Hurst (2013), we could not reproduce their results using either high-coverage data set (Fig. 6). Furthermore, upon reanalysis of the method previously employed, we found that it led to false signals of stalling in low-coverage windows—indicating apparent pausing even in simulated data where no pausing was present—and produced signals of ribosome pausing at least as strong as those observed at positive codons in regions containing no positive codons at all (Fig. 6C). In addition, the strong enrichment of positive amino acids in the 0 position of codons due to 5′ biases in read sequence in the Ingolia et al. data may also have contributed to false signals of stalling (Supplemental Fig. S18; Supplemental Material). Therefore we conclude that there is no conclusive *in vivo* evidence for a stalling effect of positive amino acids.

### Other factors associated with ribosomal stalling

Multiple studies have shown that mRNA secondary structure plays an important role in regulating translational initiation (Schauder et al. 1989; Kudla at al. 2009; Shah et al. 2013; Goodman et al. 2013). However, its importance in affecting the rate of ribosomal elongation remains controversial. For instance, a recent high-throughput analysis of secondary structure in *S. cerevisiae* reported that mRNA structure is far less extensive *in vivo* than *in vitro*, and is poorly predicted by computational methods (Rouskin et al. 2014). Furthermore, analyses of the effects of structure using yeast riboprofiling data have been inconclusive (Tuller et al. 2011; Zur and Tuller 2012; Charneski and Hurst 2013). Because mRNA structure is influenced by base content (since G:C bonds are stronger than A:U bonds), biases including the enrichment of adenines at both termini of reads overwhelms any potential signal of increased Ribo occupancy near regions of secondary structure. Therefore, riboprofiling data may not be ideal for studies of the effect of mRNA structure in the absence of methodological developments that control for biases introduced during library construction.

We observed no effect of wobble base-pairing on corrected Ribo occupancy. On the contrary, whereas no particular pattern was observed at position 5, position 6 showed a general bias towards increased occupancy of the cognate (non-wobble) codon. Therefore, the pattern of increased occupancy at G:U wobble pairs observed in nematodes and humans (Stadler and Fire 2011) does not appear to hold in yeast. Importantly however, the precise positioning of the wobble codon relative to the 5′ end of Ribo fraction reads differs between nematodes and humans, indicating that this pattern may be labile over long evolutionary distances.

### Analysis of base-level riboprofiling data

Riboprofiling data represent a significant advance over previous methods of translational analysis by enabling measurements of ribosomal occupancy across the transcriptome, without the need to experimentally perturb translational conditions in the cell. While this has dramatically increased our knowledge of genome-wide translational regulation (Ingolia et al. 2009; Ingolia et al. 2011; Li et al. 2012; Brar et al. 2012; Stadler and Fire 2013; Artieri and Fraser 2014; McManus et al. 2014), inconsistent interpretation of nucleotide-level data has produced contradictory results and made direct comparisons between studies challenging. We conclude that mitigating technical biases in riboprofiling – either experimentally or computationally – will likely reveal additional features of mRNAs that are most relevant to translational biology.

## Materials and Methods

### Riboprofiling data

The *S. cerevisiae* riboprofiling data used in this study were obtained from Artieri and Fraser (2014), McManus et al. (2014), and Ingolia et al. (2009) (Gene Expression Omnibus [GEO] entries GSE50049, GSE52119, and GSE13750, respectively). In the case of the Artieri and Fraser data, some of these samples were sequenced by multiplexing riboprofiling libraries generated from both *S. cerevisiae* and the closely related species *S. paradoxus* (Supplemental Table S3). Therefore we independently sequenced the *S. cerevisiae* Ribo fraction replicate 1 sample (deposited in NCBI Sequence Read Archive [SRA] entry SRS514738) and mapped the reads (see below) in parallel to the sample generated by sequencing the multiplexed species libraries (GEO sample GSM1278062). The strong congruence of estimated RPKMs between the individual and the multiplexed sequencing samples (rho = 0.995, p < 10^−15^; Supplemental Figure S21), as well as the biological replicates (Supplemental Fig. S1) indicated that the stringent mapping method successfully identified *S. cerevisiae* reads from the mixed sample. Supplemental Table S3 indicates the sources of the individual replicates.

### Riboprofiling library mapping

Reads from both fractions of all datasets were mapped in a strand-specific manner using the iterative method described in Ingolia (2010). We first excluded reads that mapped to the complete rDNA sequence of *S. cerevisiae* when trimmed to a length of 23 nt from the 5′ end using Bowtie version 0.12 (Langmead et al. 2009) allowing 3 mismatches and a maximum of 20 mapping locations. Remaining reads were mapped to the *S. cerevisiae* strain S288c genome (R61-1-1, 5^th^ June 2008) allowing no multimappers and no mismatches. Mapping reads were filtered such that no more than 30 bp (31 bp if the 3′ most base was an A), and no less than 27 bp (28 if the 3′ most base was an A) mapped (filtering based on terminal A bases accounts for the potentially spurious adenines added during the poly-A polymerase mediated reverse transcription priming). Only protein-coding genes with 40 or more codons were analyzed, and mapping reads were assigned to the CDS if their 5′-most base mapped at or between the 16^th^ codon and 16 codons before the end, in order to avoid effects of ribosomes paused near the start and stop codons (Ingolia et al. 2009; Ingolia et al. 2011).

The read mapping length distribution (Supplemental Fig. S3) was determined using the iterative trimming method as above on all non-rRNA mapping reads, but instead beginning with reads trimmed to 35 bp (the shortest read length generated among all three datasets) and trimming one nucleotide at a time until reaching 23 nt, retaining the longest mapping read length. Barplots were then generated by determining the percentage of reads mapping at each length among all mapping reads.

### Identifying technical biases in riboprofiling data

Mapped reads were separated into categories based on whether their 5′ ends mapped to the first, second, or third reading frame of codons. The relative proportion of each base among reads was calculated for the first 27 nucleotides of each read (corresponding to the minimum mapping read length; for Supplemental Fig. S6 the number of nucleotides analyzed was extended accordingly). Nucleotide bias was then determined by scaling the proportions of each base within each reading frame by its mean proportion across all of the same positions within codons (i.e., all first, second, or third positions) in the 27 nucleotides, thereby accounting for codon-position specific differences in expected base compositions. The ratios were log_2_ transformed for the purpose of plotting. In order to determine the degree of over-representation of adenines at the 5′ ends of reads, the proportion of adenines in the 1^st^ nucleotide position of mapping reads was compared to the proportion of adenines within the CDSs of analyzed genes (see also Supplemental Table 2).

The corresponding codon bias was determined by a fraction-specific method analogous to that presented in Fig. 3: The 5′ ends of reads from each fraction were mapped separately and the codon-level coverage was determined, retaining only codons with 5′ mapped reads in both fractions for analysis. Within each gene, codon-level coverage values for each fraction were separately scaled by the mean codon-level coverage of analyzed codons in order to account for coverage differences among genes. These scaled values were then log_2_ transformed (e.g., log_2_[scaled mRNA coverage] or log_2_[scaled Ribo coverage]) and then applied to the 5′ mapping codon and to the eight consecutive codons downstream (labeled 0-8; representing the minimum number of codons overlapped by a short read), producing a coverage value for each codon at each position. In this manner, the mean log_2_ coverage value for each of the 61 sense codons at each position was determined. We then asked whether the codons at each position were over- or under-represented relative to all nine positions by scaling the log_2_ coverage value of each codon at each position by the mean log_2_ coverage value across all nine positions – producing a new value that represents the degree to which each of the 61 sense codons deviates from its mean representation across the length of the read. To represent the degree to which each position deviated from expected codon frequencies in a graphical manner, we calculated the coefficient of variation (CV) - the standard deviation expressed as a percentage of the mean - across 61 sense codons at each position, where higher CVs indicate positions with a greater deviation from expected codon proportions.

### Determination of position-specific corrected Ribo coverage

In order to account for mapping biases shared between the mRNA and Ribo fractions in a position-specific manner, Ribo fraction occupancy was scaled by that of the mRNA fraction in the manner outlined in Fig. 3: The 5′ ends of reads from both fractions were mapped as detailed above and the codon-level coverage was determined for each fraction separately, retaining only codons with 5′ mapped reads from both fractions for analysis. Within each gene, codon-level coverage values were scaled by the mean codon-level coverage of analyzed codons. These scaled values were used to calculate the log_2_(Ribo/mRNA coverage) for each codon, accounting for shared biases between the two fractions. This log_2_(Ribo/mRNA coverage) was then applied from −8 to +8 codons relative to the codon overlapped by the 5′ end (representing 17 codons in total). Performing this analysis over all positions with data within the coding transcriptome produced a distribution of log_2_(Ribo/mRNA coverage) values for each codon at each of the 17 positions representing that codon’s contribution to ribosomal pausing, given its position relative to the ribosome-protected fragment (represented in tabular format by the mean log_2_[Ribo/mRNA coverage] of each codon at each position). The relative enrichment of each codon at each position was determined by scaling its mean log_2_(Ribo/mRNA coverage) value by the mean value of all codons at that position such that codons with positive log_2_ values were enriched relative to expectations and those with negative values were depleted. Note that in cases where a codon was not represented at a particular position, which only occurred when data were downsampled or divided into low coverage subgroups, the codon was given a log_2_(Ribo/mRNA coverage) value of 0 at that position.

As a negative control the analysis was re-run 100 times on datasets in which the genomic coordinates of the 5′ ends of mapping reads were preserved, but where the order of the codons within each gene was shuffled at random. Note that the start and stop codons were not included in the shuffling as these were always excluded in the analysis by virtue of the exclusion of codons at the beginning and ends of transcripts (see above).

### Analysis of factors implicated in affecting rates of translation

Codons were grouped into standard biochemical categories (i.e., positively charged, negatively charged, polar non-charged, and hydrophobic) plus an additional ‘special’ category containing cytosine, glycine, and proline. Wobble base positions in *S. cerevisiae* were obtained from Percudani and Ottonello (1999). Positions within mRNAs in either single-stranded or double-stranded conformation were obtained from Ouyang et al. (2013). The three different optimality measures used were relative synonymous codon usage (RSCU; Sharp and Li 1987), absolute adaptiveness (Wi; dos Reis et al. 2004) and the normalized translational efficiency scale (nTE; Pechmann and Frydman 2013).

### Application of the Charneski and Hurst method

The r_pos_/r_prec30_ values for 61 codon windows centered on the first positive amino acid encoding codon of a cluster of positive charges were determined as indicated in Charneski and Hurst (2013). The number of positive amino acids in each cluster (one, two, three, four or five, and six or more) as well as the maximum number of codons spanned by a cluster were also defined as in (Charneski and Hurst 2013). Codon level coverage was calculated as the mean nucleotide coverage within a codon. To reproduce the results of the original analysis, we combined the replicate data as per their method: calculating the nucleotide-level coverage as the average of the coverage determined in each replicate. Note however, that unlike the original analysis, we did not map reads to the mitochondrial transcriptome as it is unclear whether translational dynamics are affected by differences between the cytoplasmic and mitochondrial ribosomes and tRNA pools.

To determine whether a stalling effect was observed within regions without positive charges we identified all 61 codon windows that do not contain any positive amino acids and treated the center codon as the focal position for calculating the r_pos_/r_prec30_ values. As many such windows are immediately adjacent to one another (e.g., a run of 70 non-positive amino acids will contain 10 possible 61 codon windows), we subsampled a number positions equivalent to the number of ‘1 positive charge’ clusters used to draw panel 1 of Supplemental Fig. S17 at random 100 times from all possible windows lacking any positive amino acids, and averaged the r_pos_/r_prec30_ values over the replicate subsamples. To test whether the stalling effect of subsampled data was significantly different from the observed data, we performed Kruskal-Wallis rank sum tests (see below) on the distribution of AUC values from all of the positions analyzed in the actual data in comparison to the mean AUCs of the 100 randomly sampled replicates.

In order to explore how lower read coverage influenced the appearance of ribosomal slowing in 61-codon windows, we simulated either 10, 100, or 1000 reads per CDS with random probability in their mapping location, and equal probability of any length from 27 to 30 nt. The start and stop positions of the CDSs were based on the definition of CDS mapping reads used in Charneski and Hurst (2013): i.e. the 5′ end of reads mapped between 16 nt before the start and 14 nt before the end of the annotated CDSs.

### Statistics

All statistics were performed using R version 2.14.0 (R Core Team 2013) in addition to custom Perl scripts. 95% confidence intervals were empirically determined from the distribution of log_2_(Ribo/mRNA coverage) values from the data using the ‘boot’ package (Davison and Hinkley 2008). Kruskal-Wallis tests were performed using 10,000 permutations of the data as implemented in the ‘coin’ package (Hothorn et al. 2008).

## Acknowledgements

We thank members of the Fraser lab and the Stanford Whole Genome Sequencing group for useful comments on earlier versions of this work. This work was supported by a Natural Sciences and Engineering Research Council of Canada Postdoctoral Fellowship to CGA and NIH grant 1R01GM097171-01A1. HBF is a Sloan Fellow and Pew Scholar. The funders had no role in study design, data collection and analysis, decision to publish, or preparation of the manuscript.

## Author Contributions

CGA and HBF designed the study. CGA performed all experimental work and data analysis. CGA and HBF wrote the manuscript.

## Data Access

The raw sequencing reads generated in this study are deposited in the NCBI Sequence Read Archive under accession number SRS514738. The locations of all other data are indicated in the Materials and Methods and Supplemental Table S3.

## Supplemental Materials

### Comparison of 5′ and 3′ biases among the three datasets

The most pronounced deviations from expected nucleotide frequencies were observed at the 5′ ends of both fractions in all three datasets, though 3′ biases were also present in some cases (see below) (Fig. 2; Supplemental Figs S4–S6). The specific patterns of bias observed differed in a manner likely reflecting differences in library construction protocols used. Both the Artieri and Fraser and Ingolia datasets performed circularization of the DNA products of *E. coli* poly-A polymerase mediated reverse transcription, leading to most reads containing spurious adenine nucleotides at their 3′ ends (Artieri and Fraser 2014; Ingolia et al. 2009). Therefore, mapping was accomplished by iteratively trimming the 3′ ends and retaining the longest mapping length within the acceptable range while discarding reads that mapped at either longer or shorter lengths. As a consequence, reads tended to map with greater length if they aligned to regions of the genome that happen to harbor adenines in the 3′-most end of the mapping location, as revealed by an increasing relationship between 3′ adenine bias and read mapping length in some libraries (Supplemental Fig. S6). Unlike 5′ bias, this cannot affect the position of mapped reads, but it did increase the base-level coverage of adenine residues, which could create spurious signal in some analyses.

In contrast to the two other datasets, the McManus et al. data employed a universal miRNA linker in order to prime reverse transcription, which appears to have mitigated the 3′ adenine bias (McManus et al. 2014) (Fig. 2, Supplemental Fig. S6). However, this appears to have introduced a more strongly pronounced bias against cytosine at the 4^th^ nucleotide position as compared to the other datasets. We note that the McManus et al. protocol also differed from the two other datasets in the manner of sucrose gradient mediated isolation of the 80s monosome, as well as in the specific details of the size-selection step subsequent to the addition of the miRNA linker. These may explain the more restricted mapping length distribution we observed in the 28 – 31 nt range in the this dataset (Supplemental Fig. S3).

Despite differences among protocols, a preference for adenine in the first nucleotide position remained a consistent bias among all datasets, indicating that some component of the library preparation protocol common to both fractions preferentially selected for certain fragments. However, the ‘non-preferred’ nucleotides varied among datasets – cytosine and guanine in the Artieri and Fraser and McManus et al. data vs. guanine and thymine in the Ingolia et al. data – which could reflect changes in next-generation sequencing protocols, reagents, and/or equipment used: The Ingolia et al. dataset was generated on the Illumina Genome Analyzer II instrument using reagents available at the time, whereas the two more recent datasets were generated using the Illumina HiSeq 2000 instrument.

### Analysis of nucleotide and codon biases based on mapping read length

Previous studies have identified reads mapping at a length of 28 nt as being of particularly ‘high-quality’ as this corresponds to the fragment length that the ribosome should protect from nuclease digestion during library preparation (e.g., Ingolia et al. 2009; Qian et al. 2012). We therefore reanalyzed reads as in Fig. 2, but this time as a function of mapping length (Supplemental Fig. S6). While mapping read length had no substantial effect on biases in the mRNA fractions, 28 nt mapping reads from the Ribo fractions of all three datasets showed the strongest internal biases in codon representation (though internal enrichment of 27 nt reads were similar). Furthermore, 28 nt reads were enriched among first reading frame mappers in all three datasets, whereas third frame mappers were enriched for longer read lengths (Supplemental Fig. S7). Hence, if internal biases in codon representation represent a signal of ribosome stalling, reads ≤ 28 nt appeared to be optimal for detecting such a phenomenon.

### Applying the corrected Ribo coverage method to the data of Ingolia et al.

When the corrected Ribo coverage method was applied to the two datasets generated by Ingolia et al., we did not observe a consistent enrichment of proline codons at position 4 (the P-site), in contrast to the higher-coverage datasets (Supplemental Fig. S11). Furthermore, we could not attribute this difference to the lower coverage of these data as subsamples of first frame mappers of the Artieri and Fraser or McManus et al. data to the level of coverage of the Ingolia et al. rich dataset continued to produce strong enrichment of the four proline codons (Supplemental Fig. S11). However, we note that features unique to the Ingolia et al. dataset render it qualitatively different from the other two sets. First, stronger differences in 5′ nucleotide and codon bias were observed between the two biological replicates of the Ingolia et al. data as compared to the other two sets, suggesting the presence of substantial batch effects across library preparation and/or sequencing (Supplemental Fig. S5). Second, in addition to these differences, the magnitude of 5′ biases in all Ingolia et al. Ribo fractions was substantially higher than their corresponding mRNA fractions, indicating that the Ribo fractions harbored more significant 5′ technical biases that could not be controlled by those observed in the mRNA fraction (in comparison, the 5′ biases are of comparable magnitude between fractions of the Artieri and Fraser and McManus et al. datasets) (Fig. 2, Supplemental Figs. S4–S6).

We argue that these biases are technical and not biological in nature because they were highly specific to the 5′ ends of Ribo reads and were not observed at adjacent nucleotide/codons sites, either upstream or downstream, as would be expected if, for instance, previously translated amino acids were hindering ribosomal progression due to interactions with the negatively charged exit tunnel (Lu et al. 2007) (see Supplemental Fig. S18). Therefore, we suggest that strong patterns of bias observed at the 5′ ends of the Ribo reads in the Ingolia et al. data – as compared to the Artieri and Fraser or McManus et al. data – may have led to restricted patterns of mapping that interfered with the ability to detect patterns of over-represented codons at other sites.

### Analysis of RNA secondary structure

Previous studies reported a correlation between ribosomal occupancy of the Ingolia et al. data and the presence of secondary structure in mRNAs (Tuller et al. 2011; Charneski and Hurst 2013). Therefore we determined whether such a relationship was observed in the Artieri and Fraser and McManus et al. datasets. We obtained experimentally determined structures for 2,839 *S. cerevisiae* mRNAs (Ouyang et al. 2013) and tested whether the local secondary structure (from 10 codons upstream to 20 codons downstream of the read) was correlated with coverage of the 5′ ends in the Ribo, mRNA, or corrected Ribo reads (Supplemental Fig. S13; See Supplemental Methods).

We observed that the overall correlation between read occupancy and secondary structure was weak at all positions (Supplemental Fig. S13). However, the strongest correlations were observed at codons corresponding to the 5′ and 3′ ends of reads, coinciding with the locations of the most pronounced nucleotide biases (Fig. 2). For example, in the correlation with the mRNA fraction of the Artieri and Fraser data, secondary structure at codons corresponding to the ends of reads were negatively correlated with occupancy, as expected from the over-representation of adenine-rich codons at these positions, which do not form strong base-pairing interactions. Despite a preference for adenines at the 3′ ends of reads, the Ribo fraction showed a slight increase in the correlation between secondary structure at codon position 9 and read occupancy (Supplemental Fig. S13); however, it remained weaker than the correlations observed in the mRNA fraction. The McManus et al. data showed the strongest correlations at the 5′ ends, also consistent with terminal nucleotide biases dominating the signal. Therefore, our results suggest that if there is a correlation between ribosomal occupancy and the presence of secondary structure, it is quite weak relative to the general variation in ribosomal occupancy across mRNAs – so much so that the signal is overwhelmed by the inherent biases introduced during the library construction process (see Discussion of main text).

### The rate of translation is not correlated with codon optimality

We failed to find a significant negative correlation between corrected Ribo occupancy and any of three different measures of codon optimality at either position 4 (Supplemental Fig. S14) or position 5 (Supplemental Fig. S15). This agreed with other riboprofiling-based analyses performed in bacteria (Li et al. 2012), yeast (Qian et al. 2012; Zinshteyn and Gilbert 2013), and mouse (Ingolia et al. 2011). In contrast, those studies that reported such a relationship focused primarily on the association between non-optimal codons and increased ribosomal occupancy at the 5′ ends of genes (Tuller et al. 2010b; 2011), rather than a direct association between Ribo fraction occupancy and non-optimal codons. In addition, a recent systematic study of translation in *E. coli* using synthetically designed N-terminal codon compositions found that the preference for non-optimal codons near start codons could be explained by their reduced secondary structure, enabling efficient translational initiation (Goodman et al. 2013). Therefore, our analysis supports the notion that the pool of tRNAs and transcriptome-wide CUB are adapted for efficient peptide synthesis *in vivo*, and that previous *in vitro* studies that found a strong relationship between CUB and translation rate may reflect circumstances that deviate significantly from those found in the cell (Plotkin and Kudla 2011; Qian et al. 2012).

### Reanalysis of the approach of Charneski and Hurst

To illustrate the read coverage dependence of the method of Charneski and Hurst (2013), consider a situation where data are extremely sparse, such that no more than a single read maps to any 61 codon window (which is the size of the analysis space employed in their manuscript, representing 30 codons upstream of a putative stalling codon, to 30 codons downstream; Supplemental Fig. S19). Three possible mapped reads are shown to illustrate that read mapping position strongly influences its contribution to the average r_pos_/r_prec30_ value. Importantly, averaging over all possible single read mapping positions produces a pattern that is highly characteristic of what is interpreted as the typical ‘stalling’ pattern observed in several of Charneski and Hurst’s figures. This bias towards producing a signature of stalling can be further illustrated by generating randomly positioned reads within the analyzed annotation, with length randomly chosen between 27 and 30 nt, and with increasing levels of coverage. When averaging over all possible 61 codon windows, a stalling pattern very similar to those observed in the Charneski and Hurst analysis manifests at low read coverage, but largely disappears at high coverage (Supplemental Fig. S19). Tellingly, averaging over all possible codon sites in the analyzed data produces a pattern very similar to that observed from 100 random reads assigned to each gene (note that the average read depth of the data is ~260 reads per gene and the reads are not randomly distributed as in the simulated data). It is also worth noting that the biases observed in this approach also explain the consistent pattern of increased coverage at the left end (near the −30 position) of almost all figures in the Charneski and Hurst manuscript, which the authors attribute to “some residual slowing […] due to slowing elements (e.g., positive charges) that may be encoded just upstream…” (pg. 4). It is clear from simulated data that this pattern results from the edge effect of low coverage reads that partially overlap the most upstream codons of the window being considered.

We also note that the most pronounced codon level biases observed in the Ingolia et al. Ribo fractions are an enrichment of positive amino acid encoding codons at the 5′ end of reads, likely the result of the most abundant positive charge encoding codons being A rich (particularly among first frame mappers): lysine, AA[A/G], and arginine, AG[A/G] (Supplemental Fig. S18). Irrespective of the coverage sensitivity of the r_pos_/r_prec30_ method, this bias could explain why the Ingolia et al. data produce an accumulation of reads when positive amino acid encoding codons are at and downstream of the focal codon.

## Supplemental Methods

### Analysis of mRNA secondary structure

We obtained the data of Ouyang et al. (2013), which includes binary designations for each base in 2,839 of the mRNA transcripts analyzed in this study, identifying it as single or double-stranded according to the transcriptome-wide measurements of Kertesz et al. (2010). Using the codon-specific occupancy for the 5′ end of first position mapping reads, we determined the correlation between occupancy at codon position 0, corresponding to the 5′ end of the read, and secondary structure from codon position −10 to +20. Codon-level secondary structure was scored as the average structural value of the three nucleotides in each codon, where a single-stranded nucleotide was given a value of 0 and a double-stranded nucleotide, 1. This correlation was determined independently using the mRNA and Ribo fractions in addition to the corrected Ribo coverage in order to determine whether biases in either fraction drove any patterns observed in the corrected data.

### Analysis of Ribo fraction codon-level biases

In order to generate Supplemental Fig. S18, we followed the method to generate corrected Ribo coverage, but without correction by the mRNA fraction: The 5′ ends of reads from the Ribo fraction were mapped as detailed in the methods section and the codon-level coverage was determined, retaining only codons with 5′ mapping data for analysis. Within each gene, codon-level coverage values were scaled by the mean codon-level coverage of analyzed codons in order to account for coverage differences among genes. These scaled values were then log_2_ transformed (e.g., log_2_[scaled Ribo coverage]) and then applied from −8 to +8 codons relative to the codon overlapped by the 5′ end (representing 17 codons in total). Performing this analysis over all positions with data within the coding transcriptome produced a distribution of log_2_(scaled Ribo coverage) values for each codon at each of the 17 positions, which were then combined into biochemical categories. The relative enrichment of each category at each position was determined by scaling its mean log_2_(scaled Ribo coverage) value by the mean value of the five categories at that position such that categories with positive log_2_ values were enriched relative to expectations and those with negative values were depleted. Error bars were calculated as the standard error of the mean among all measurements of codons within a biochemical category at each position.

**Supplemental Table S1.** Number of reads in each dataset mapping to the *S. cerevisiae* annotation used in the current analysis.

| Sample | Fraction | Replicate | Mapped Reads |
| --- | --- | --- | --- |
| Artieri and Fraser | Ribo | 1 | 26,648,753 |
|  |  | 2 | 17,102,202 |
|  | mRNA | 1 | 10,128,586 |
|  |  | 2 | 21,137,349 |
| McManus et al. | Ribo | 1 | 6,246,282 |
|  |  | 2 | 7,580,892 |
|  | mRNA | 1 | 2,153,414 |
|  |  | 2 | 2,050,551 |
| Ingolia et al. rich | Ribo | 1 | 711,601 |
|  |  | 2 | 958,424 |
|  | mRNA | 1 | 288,061 |
|  |  | 2 | 1,340,197 |
| Ingolia et al. starved | Ribo | 1 | 350,787 |
|  |  | 2 | 724,985 |
|  | mRNA | 1 | 127,623 |
|  |  | 2 | 796,049 |

**Supplemental Table S2.** Proportion of adenine in the CDS as a function reading frame and expression level. The proportion of adenine within the CDS presented in the main text is calculated as the proportion among all nucleotides. As more highly expressed genes show greater CUB, this value will vary based on the subset of genes analyzed as well as the reading frame within each codon. However, the proportion of adenine varies within a narrow range, especially among first frame mappers. Expression quartiles were determined based on the mean RPKM among both replicates of the Artieri and Fraser data.

| Expression Quartile | Reading Frame | Percent Adenine |
| --- | --- | --- |
| 1 | 1 | 33.5 |
|  | 2 | 33.1 |
|  | 3 | 29.3 |
| 2 | 1 | 34.3 |
|  | 2 | 36.1 |
|  | 3 | 30.1 |
| 3 | 1 | 33.3 |
|  | 2 | 35.9 |
|  | 3 | 29.4 |
| 4 | 1 | 31.5 |
|  | 2 | 35 |
|  | 3 | 26.9 |

**Supplemental Table S3.** SRA sample ID numbers for Artieri and Fraser (2014) data used in the analysis.

| Sample | Source | SRA Sample ID |
| --- | --- | --- |
| mRNA Rep. 1 | Artieri and Fraser mixed *S. cerevisiae*/*S. paradoxus* sample | SRS509272 |
| mRNA Rep. 2 | Artieri and Fraser *S. cerevisiae* sample | SRS469853 |
| Ribo Rep. 1 | *S. cerevisiae* sample generated for the present study | SRS514738 |
| Ribo Rep. 2 | Artieri and Fraser mixed *S. cerevisiae*/*S. paradoxus* sample | SRS509342<br>SRS509345 |

**Supplemental Figure S1.**
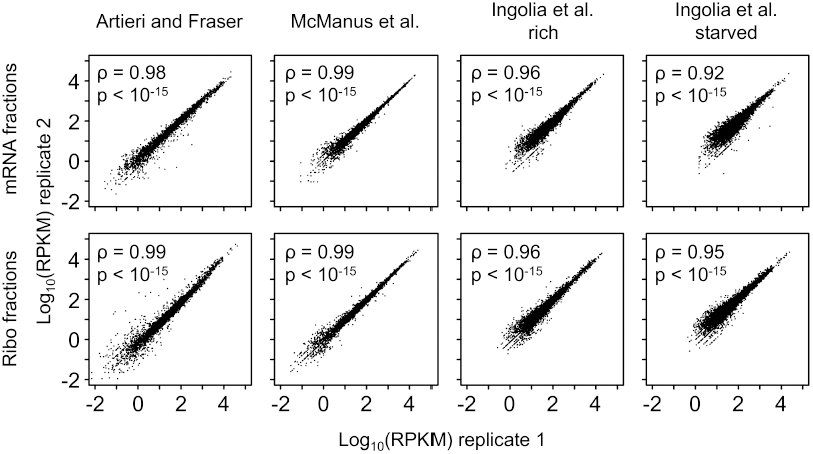
Inter-replicate correlation of expression level estimates for the analyzed datasets. Only genes with <u>R</u>eads <u>P</u>er <u>K</u>ilobase per <u>M</u>illion mapped reads (RPKM) > 0 are plotted. mRNA fractions are plotted in the row above, while Ribo fractions are indicated below. Spearman‘s ρ and associated p values are indicated in each panel. The lower ρ values in the Ingolia et al. data likely reflect the lower number of mapping reads in that dataset (Table S1). Note that in the original Ingolia et al. (2009) analysis, correlations were calculated using only genes with ≥ 128 mapping reads, which improves correlation coefficients.

**Supplemental Figure S2.**
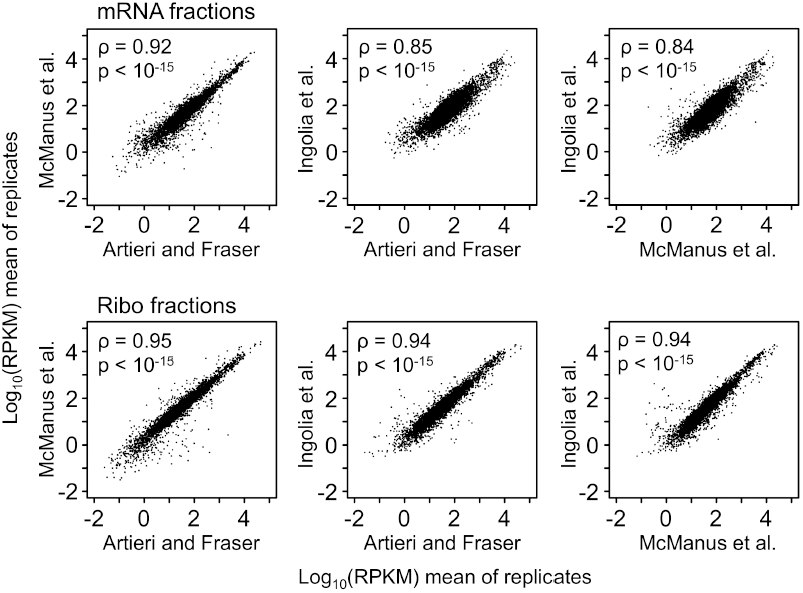
Correlation of expression levels between the datasets used in the study. The mean RPKM of the two replicates generated in each study is plotted along with Spearman correlation coefficients, ρ, and associated p values. The slightly lower coefficient of correlation for the mRNA data may reflect use of different methods to extract mRNA from the raw lysate (Ingolia et al. 2009; Artieri and Fraser 2014; McManus et al. 2014).

**Supplemental Figure S3.**
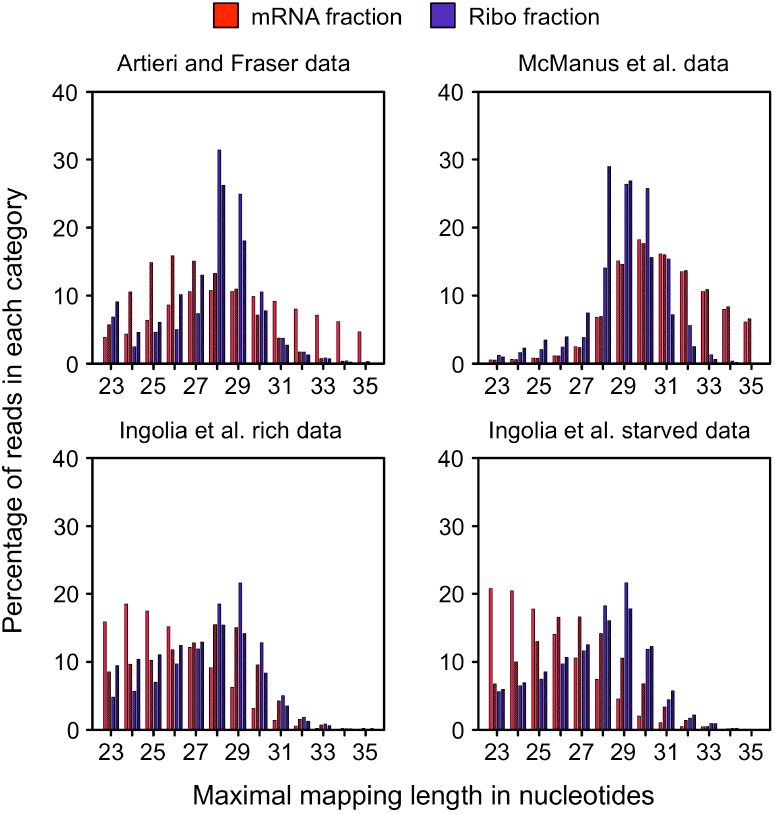
Read mapping length distribution for each of the analyzed datasets. Variation in the length distribution of the mRNA fraction (red) is likely driven by the size and precision of the fragment excised from the denaturing SDS-PAGE gel during library construction (Ingolia 2010). In contrast, enrichment of ~28 nt fragments is expected in the Ribo fraction (blue) as this is the length of mRNA occupied by a translating ribosome (Ingolia et al. 2009). Biological replicates are plotted next to one another where replicate 1 is plotted without hashes and replicate 2 is hashed, resulting in a darker shade. The distribution of the McManus et al. data is more compact and excludes short-length reads to a greater degree than other datasets. This is possibly due to their unique use of universal miRNA linkers followed by a modified size-selection protocol (see above) (McManus et al. 2014).

**Supplemental Figure S4.**
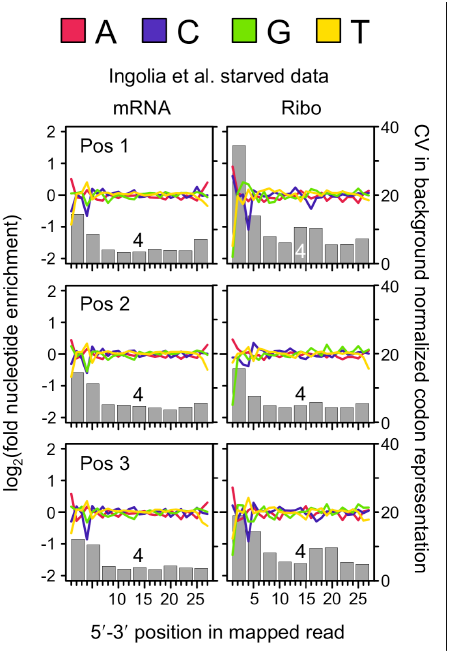
Reproduction of Fig. 2 showing that the Ingolia et al. starved data are qualitatively similar to the rich data.

**Supplemental Figure S5.**
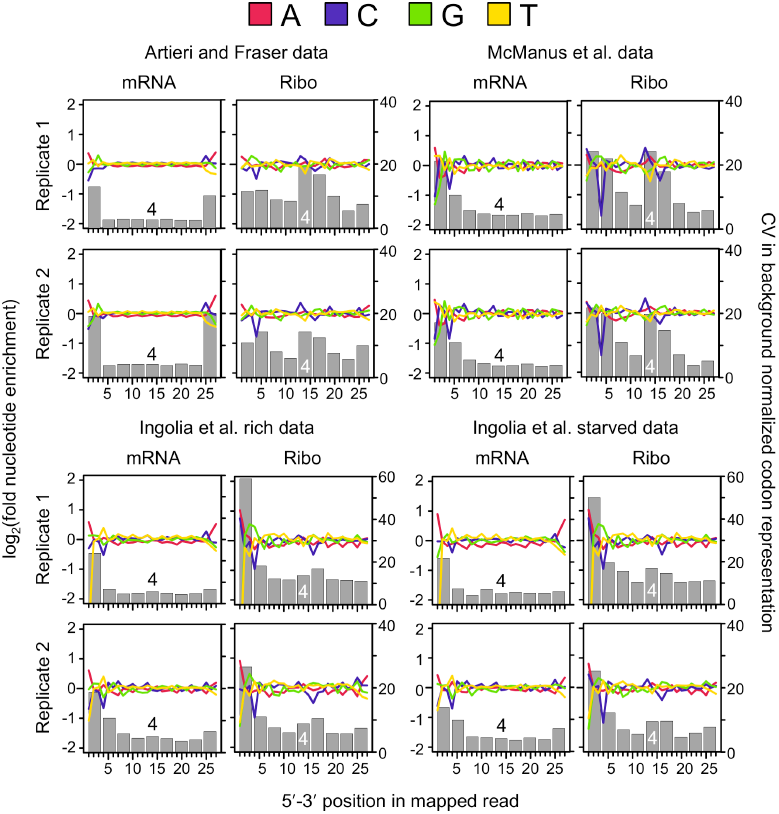
Comparison of patterns of bias across replicates for first codon position mappers. Patterns of bias were consistent across replicates of both fractions of the Artieri and Fraser and McManus et al. data (top). In the Ingolia et al. data (bottom), 5′ biases were much more pronounced in the first as compared to the second replicate (note the difference in scale for the CV in background normalized codon representation used in replicate 1). In addition to differences between replicates in the Ingolia et al. data, patterns of 5′ bias among the Ribo fractions were of greater magnitude relative to the mRNA fractions as compared to the higher-coverage datasets.

**Supplemental Figure S6.**
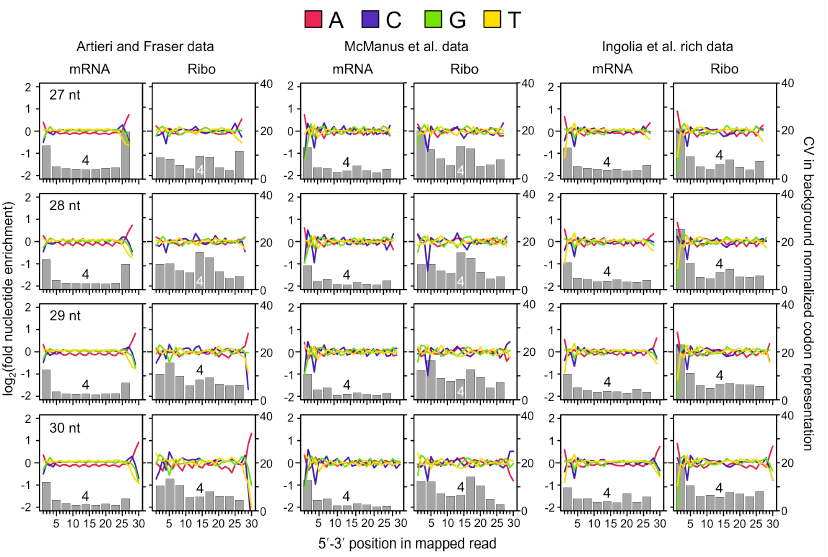
Reanalysis of patterns of nucleotide and codon bias among reads binned by mapping length. The grey bars are plotted for the 9 codon positions (0-8) corresponding to 27 nt as in Fig. 2, with the fourth codon position indicated for reference. Five prime patterns of bias are similar within fractions across mapping lengths. However, 3′ codon bias decreases with read length in the Artieri and Fraser and Ingolia et al. data due to biases in adenine content overlapping codon position 8 as a result of the use of poly-A polymerase to prime reverse transcription (see above). Significantly, in all three datasets, the strongest degree of internal codon bias in the Ribo fraction is observed among 28 nt mapping reads (with 27 nt reads often showing a similar pattern). This may be related to the general enrichment of first frame mapping reads among 28 nt mappers (Supplemental Fig. S7).

**Supplemental Figure S7.**
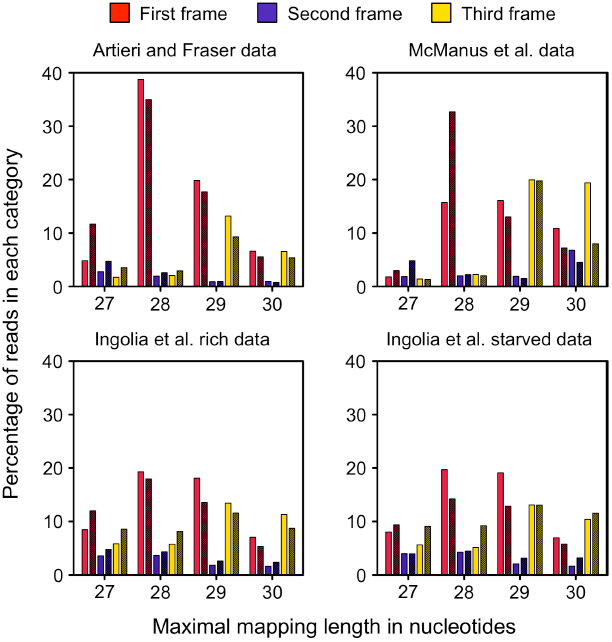
Ribo fraction reads mapping at different lengths show differential enrichment among first, second and third frame mappers. All three fractions show an enrichment of 28-29 nt mapping reads among first frame mappers, which make up the majority of reads. Third frame mappers tend to be enriched for 29-30 nt mapping reads. Few reads map to the second reading frame in any dataset. The first, second, and third reading frames are shown in red, blue and yellow, respectively. Biological replicates are plotted next to one another where replicate 1 is plotted without hashes and replicate 2 is hashed, resulting in a darker shade.

**Supplemental Figure S8.**
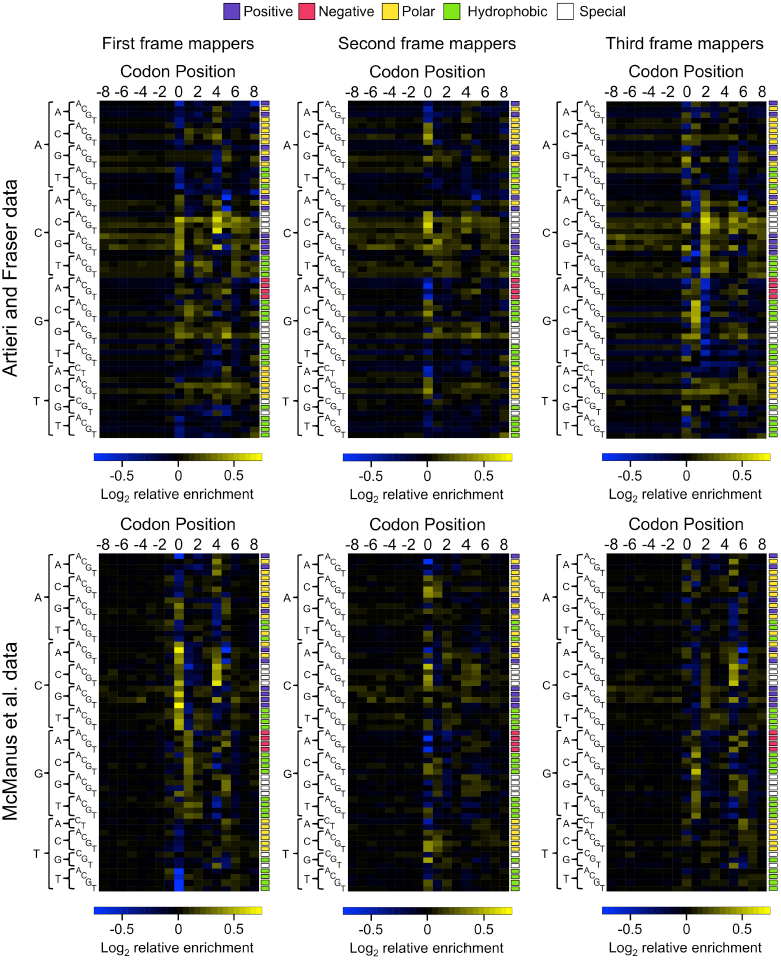
Patterns of corrected codon enrichment at positions −8 to 8 for first, second, and third reading frame mapping reads for the Artieri and Fraser and McManus et al. data. All three frames continue to show substantial bias at codon position 0, corresponding to the 5′ end of the read, indicating that despite controlling for shared biases between fractions, substantial fraction-specific 5′ end biases remain. However, first position mappers of both datasets show a clear pattern of internal enrichment at position 4, previously defined as corresponding to the ribosomal P-site. Second frame mappers show a weaker pattern of enrichment corresponding to the same codons, but it is distributed among positions 4 and 5, suggesting that precise location of a potentially active ribosomal site is less clearly defined in relation to the 5′ end among reads mapping in this frame. Similarly, reads mapping to the third frame also show a qualitatively similar pattern of codon enrichment to first frame reads, but the most highly enriched codon position is 5. This supports our observation that codon positions of third frame mappers tend to be offset by +1 codon (see main text; Fig. 2). Third frame mapping reads also show stronger patterns of bias among codon positions 1 and 2, which are qualitatively consistent among the two datasets.

**Supplemental Figure S9.**
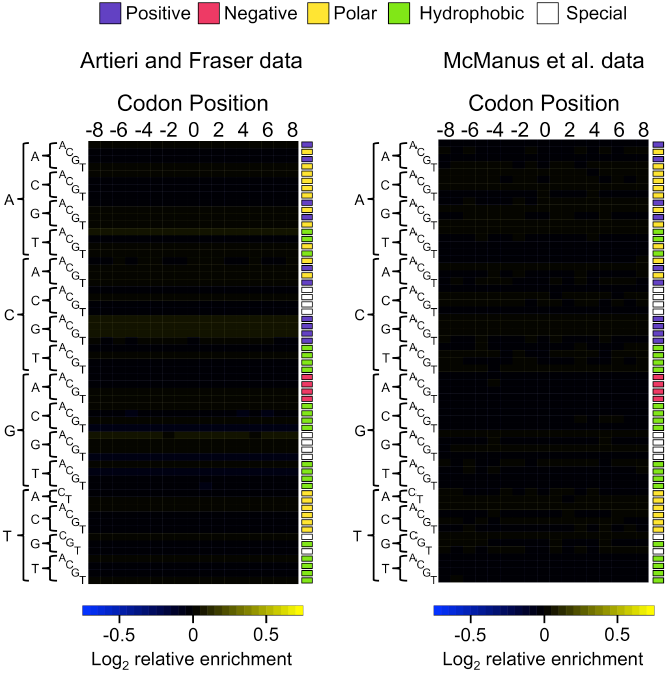
Figure 4A redrawn from the mean of 100 permutations of the codon positions within transcripts while maintaining read mapping positions. Permutations are shown using either the Artieri and Fraser or McManus et al. data. All patterns observed in the actual data disappeared. The remaining very weak enrichment/depletion represents the underlying bias in the dataset that would be observed regardless of the biological factors influencing ribosomal occupancy.

**Supplemental Figure S10.**
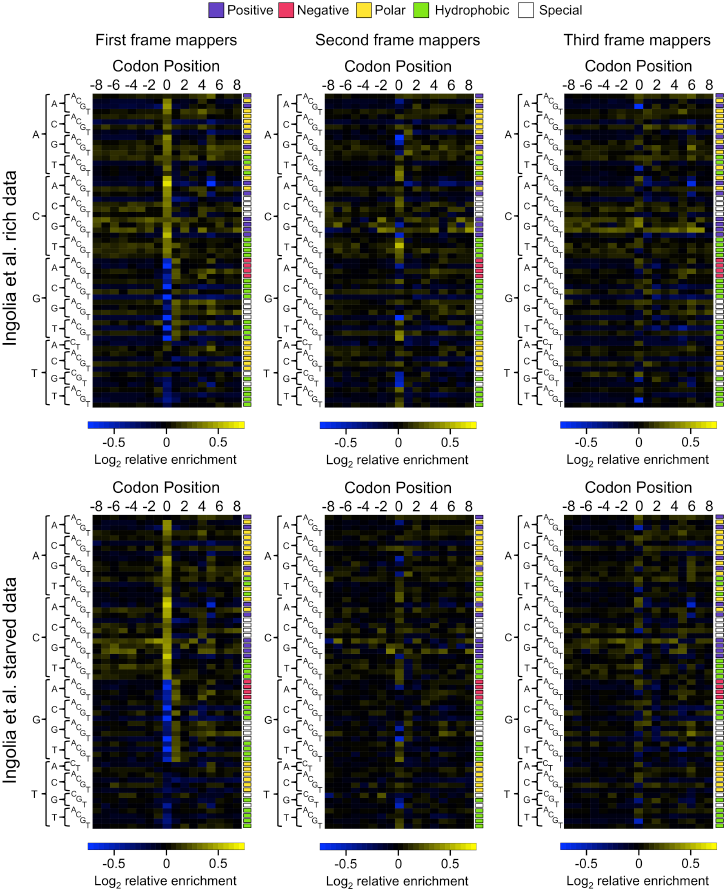
Patterns of corrected codon enrichment at positions −8 to 8 for first, second, and third reading frame mapping reads for the Ingolia et al. data. Patterns of enrichment in the Ingolia et al. data (rich, above; starved, below) differ substantially from those observed in the higher coverage datasets. For instance, first frame mappers show overwhelming codon bias at position 0 (codons beginning with A or C are universally enriched, while those beginning with G and T are depleted) whereas no upstream, nor downstream positions show strong biases as was the case for position 4 in the Artieri and Fraser and McManus et al. data. Second and third frame mapping reads also show relatively high bias at position 0 relative to other codon positions; however, their magnitude appears to be much smaller, likely due to noise introduced by the small overall number of reads mapping in these frames (Supplemental Fig. S7).

**Supplemental Figure S11.**
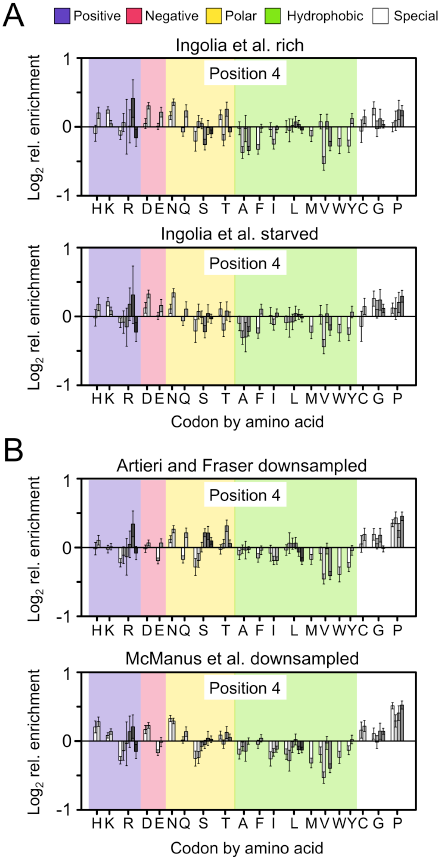
Bar plots indicating the log_2_ enrichment values at position 4 of first frame mappers. A) combined replicates of the Ingolia et al. datasets and B) Artieri and Fraser and McManus et al. datasets downsampled to the same number of reads as used to generate the Ingolia et al. rich panel. Codons are organized by amino acid using single-letter designations below and grouped by biochemical type as indicated at the top of the panel. Individual codons for each amino acid are in alphabetical order. 95% confidence intervals around the scaled enrichment values are indicated at the top of each bar. Though the Ingolia et al. data showed enrichment/depletion at similar codons to the higher coverage datasets, we did not observe the pronounced enrichment of proline (P) codons. However, this was not explained by the reduced coverage of the Ingolia et al. data, as subsamples of the Artieri and Fraser and McManus et al. data still showed a clear enrichment of all proline encoding codons (see discussion above).

**Supplemental Figure S12.**
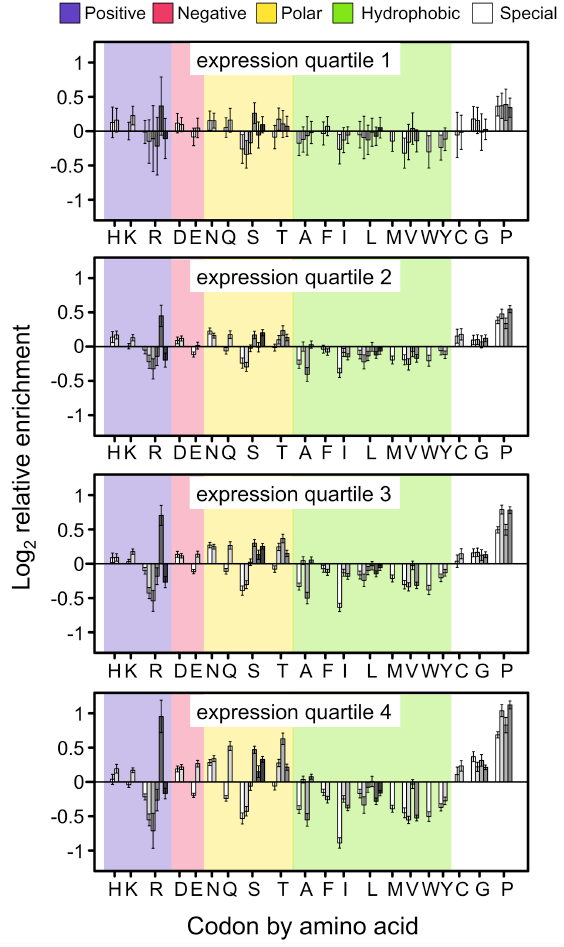
Patterns of codon enrichment at position 4 are consistent across expression levels. Genes were divided into expression quartiles based on the inter-replicate mean RPKM estimated from the mRNA fraction of the Artieri and Fraser data. Fewer codons are covered by reads in low expression genes leading to larger 95% confidence intervals, yet all proline codons remain enriched at all four quartiles. Results are qualitatively similar in the McManus et al. dataset (not shown).

**Supplemental Figure S13.**
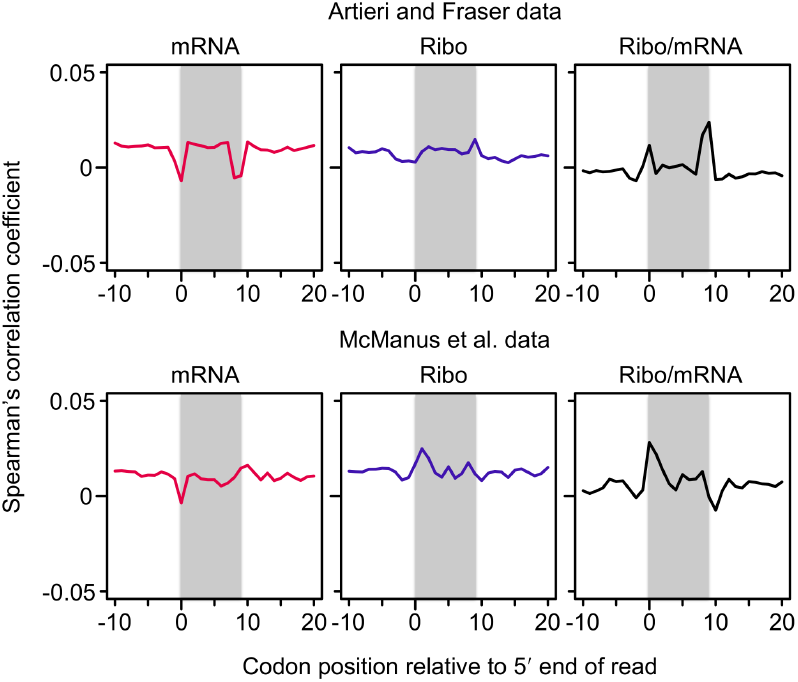
Spearman’s correlation coefficients between mRNA secondary structure and 5′ mapping codon coverage. Correlations are shown for the mRNA and Ribo fractions, as well as the corrected Ribo coverage (Ribo/mRNA). The presence of secondary structure at each codon from 10 codons upstream to 20 codons downstream was independently correlated with occupancy at the 5′ mapping codon (position 0). The grey shading indicates the 9 codons completely overlapped by each read. The overall correlation between read occupancy and secondary structure was weak at all positions in both datasets. However, it was clear that terminal nucleotide biases, particularly 5′ over-representation of adenine, which is disfavored in terms of forming secondary structure, overwhelmed other signals. In the Artieri and Fraser data, terminal adenine biases at both ends contributed to the positive correlations at positions 0 and 9 in the corrected Ribo coverage panel. The McManus et al. data lacked the 3′ adenine bias contributing to an increased positive correlation only at the 5′ end of reads. This was unlikely to be caused by secondary structure hindering ribosome progression since the 5′ ends of reads represent the trailing edge of the ribosome footprint; secondary structure would be more likely to exert an effect at the leading edge.

**Supplemental Figure S14.**
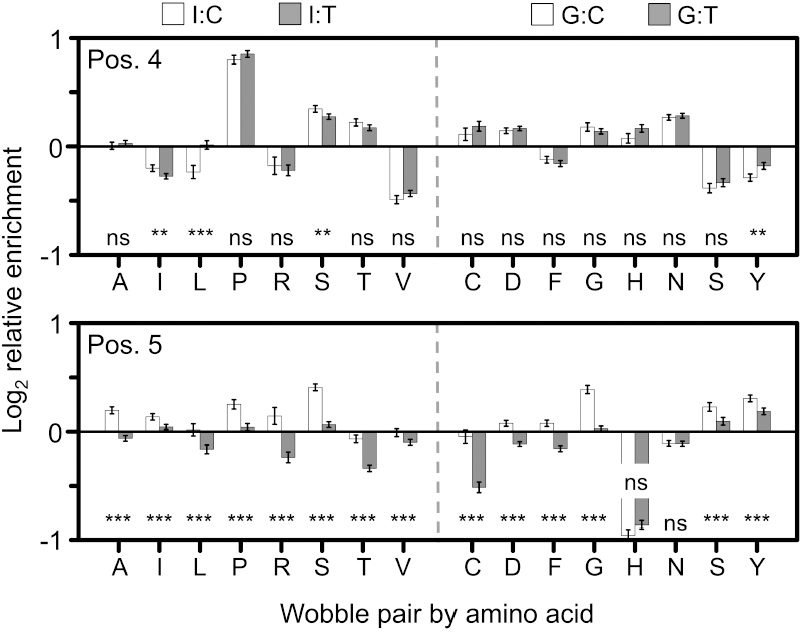
Comparison of enrichment between Watson-Crick (I:C/G:C, white) and wobble pairing (I:U/G:U, grey) codon-pairs. Amino acids using inosine (I, left) or guanine (G, right) wobble pairing are indicated. The significance of a Kruskal-Wallis rank sum test of the difference between enrichment of Watson-Crick and wobble codons is shown above each amino acid if p < 0.05 after Bonferroni correction for multiple tests (*, p < 0.05; **, p < 0.01; ***, p < 0.001). At position 4 (Pos. 4, top) there appeared to be an inconsistent general increase in enrichment of wobble pairing geometry, particularly in the case of the McManus et al data. However, at position 5 (Pos. 5, bottom) in all cases where there was a significant difference in the enrichment, Watson-Crick pairing was favored. Therefore, there did not appear to be a consistent favoring of either geometry. Furthermore, the relative enrichment differences between pairing geometries appeared to be overwhelmed by variation at individual codons themselves, irrespective of whether they use Watson-Crick or wobble pairing (e.g., Fig. 4A).

**Supplemental Figure S15.**
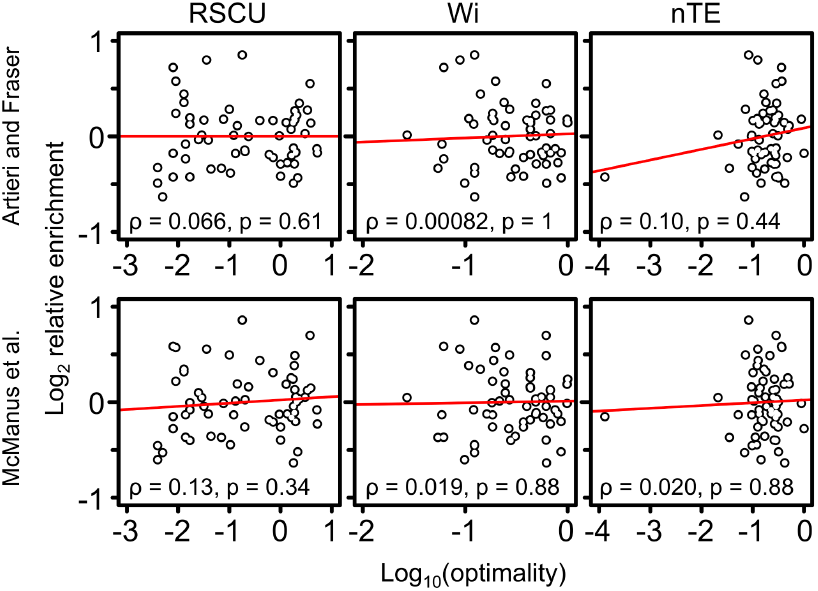
No significant correlation among three different measures of codon optimality and log_2_ corrected Ribo enrichment at codon position 4 (P-site). Spearman correlation coefficents and associated p-values are shown in each box. A negative correlation would be expected if non-optimal codons slow ribosomes. RSCU, relative synonymous codon usage (Sharp and Li 1987); Wi, absolute adaptiveness (dos Reis et al. 2004); nTE, normalized translational efficiency scale (Pechman and Frydman 2013).

**Supplemental Figure S16.**
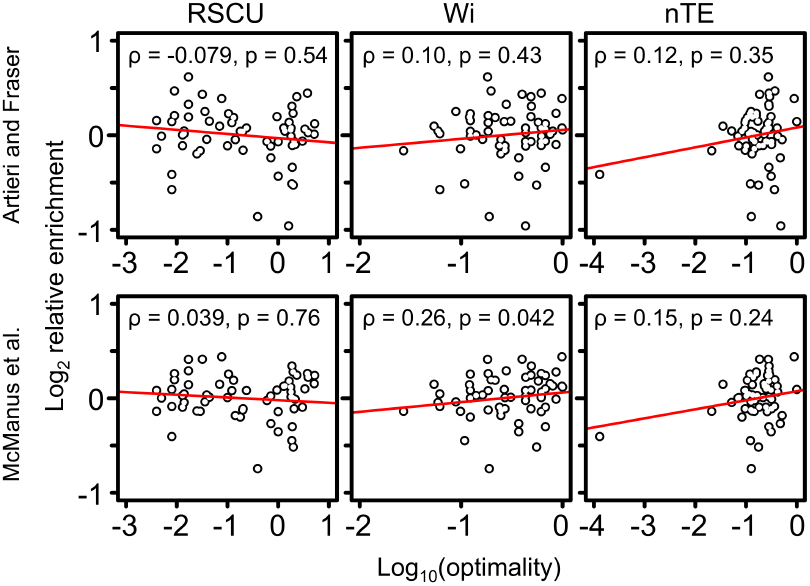
No significant correlation among three different measures of codon optimality and log_2_ corrected Ribo enrichment at codon position 5 (A-site). Spearman correlation coefficents and associated p-values are shown in each box. A negative correlation would be expected if non-optimal codons slow ribosomes. RSCU, relative synonymous codon usage (Sharp and Li 1987); Wi, absolute adaptiveness (dos Reis et al. 2004); nTE, normalized translational efficiency scale (Pechman and Frydman 2013). Note that the correlation between relative enrichment and Wi for the McManus data (p = 0.042) is no longer significant after correction for multiple tests.

**Supplemental Figure S17.**
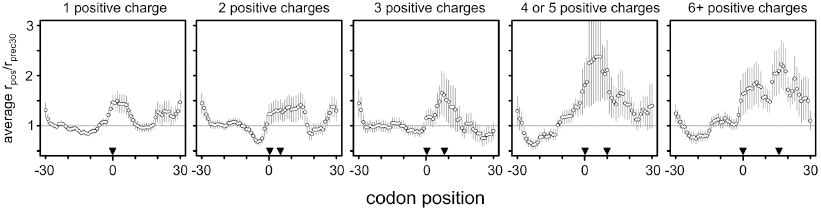
Reproduction of the additive stalling effect observed in Figure 5 of Charneski and Hurst (2013) confirming that the same analysis method was used in the present study.

**Supplemental Figure S18.**
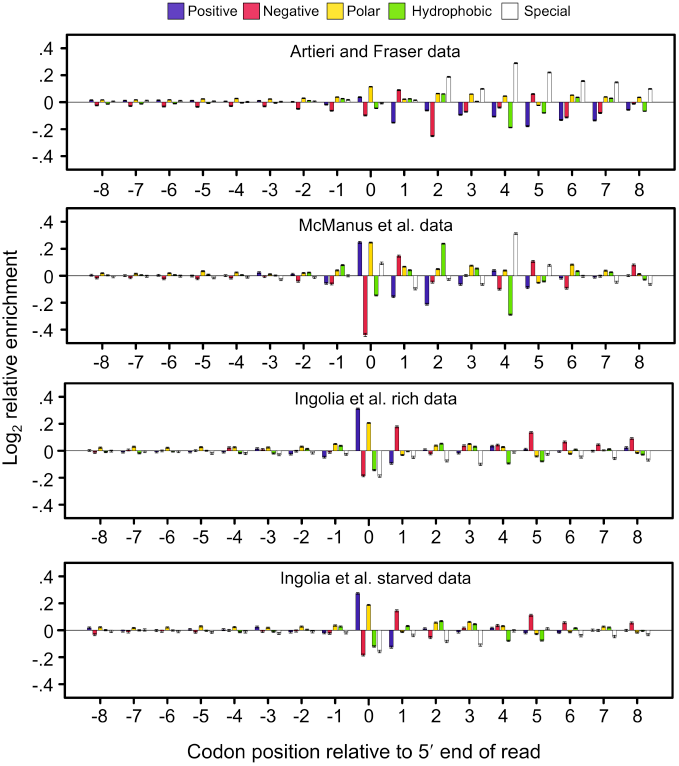
Positive amino acids are not enriched among upstream codons in the uncorrected Ribo fractions in any of the datasets, as would be expected if ribosomes were slowed as these codons passed through the exit tunnel. Enrichment was determined at the level of biochemical class using the Ribo fraction alone without correcting by the mRNA fraction (see Supplemental Methods). Reads from all read lengths and reading frames were used as in the analysis of Charneski and Hurst (2013). Error bars indicate the standard error of the mean. Biochemical classes are indicated above. Positive amino acids were not enriched in positions −8 to −1 in any dataset. However, the strongest levels of enrichment in both Ingolia et al. datasets are among positive amino acid encoding codons at position 0, which could lead to enrichment at and downstream of the focal codon used by Charneski and Hurst (Fig. 5; see above). The McManus et al. dataset also showed enrichment among positive amino acids at position 0; however, this is not as strong as the enrichment of special codons at position 4, of which proline is a member.

**Supplemental Figure S19.**
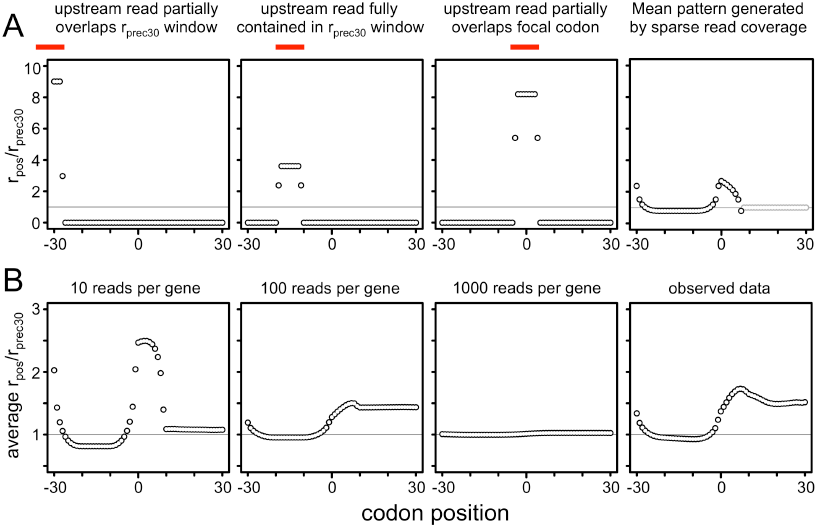
The r_pos_/r_prec30_ method of Charneski and Hurst (2013) is biased towards producing false signals of stalling when read coverage within windows is sparse. (A) When only a single read overlaps the 61 codon window of investigation, its position (indicated by the red bar) influences its contribution to the overall mean r_pos_/r_prec30_ value. Note that codon level coverage was calculated as the average coverage of its three nucleotides, allowing codons overlapping the ends of reads to have fractional coverage. Averaging over all possible 28 nt single read positions produced a characteristic ‘saddle’ pattern of stalling in the absence of any such an effect. Note that codon positions 8 to 30 are greyed out as windows containing single reads spanning them will produce r_prec30_ values of 0 leaving their r_pos_/r_prec30_ values undefined. (B) Generating randomly positioned reads of length randomly chosen between 27 and 30 nt and averaging over all possible 61 codon windows produced signals of strong stalling that only disappeared at high coverage. Each gene was assigned the indicated number of reads. The observed data also showed an overall mean pattern of stalling when averaging over all available positions. Note that the observed data have a mean coverage of ~260 reads per gene; however, this coverage is not evenly distributed as in the simulated data, and therefore does not show a pattern intermediate between that of 100 and 1000 reads per gene.

**Supplemental Figure S20.**
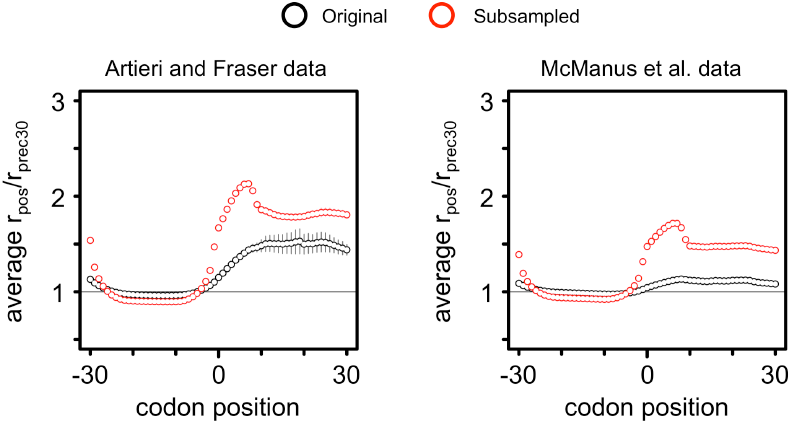
Downsampling the Artieri and Fraser or the McManus et al. data increases the ‘stalling’ effect detected. Mapped reads from both replicates of the two datasets were randomly sampled down to the mean number of reads mapping among replicates of the Ingolia et al. rich data. The average r_pos_/r_prec30_ was determined by over all 61 codon windows. In order to compare the same data directly, only sites that had mapping reads in both the original data and the subsampled data were used for plotting (black, original data; red, subsampled data). The average stalling pattern increases substantially in both datasets, highlighting the sensitivity of the method to sparse coverage.

**Supplemental Figure S21.**
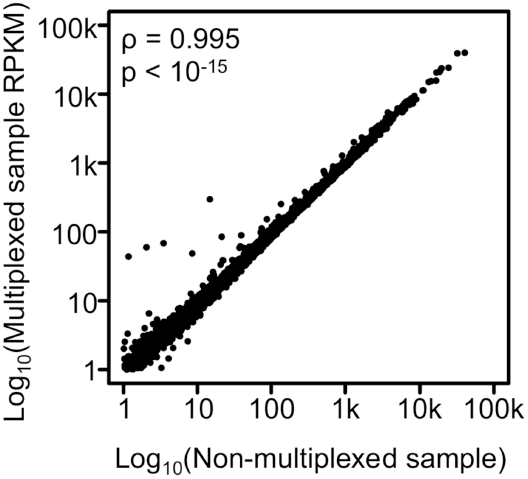
Our mapping approach identified *S. cerevisiae* reads in the multiplexed high-coverage data of Artieri and Fraser (2014). The high correlation between expression levels (in RPKM) of *S. cerevisiae* genes in the *S. cerevisiae* + *S. paradoxus* multiplexed Ribo fraction sample and the *S. cerevisiae* only sample indicated that our mapping method robustly identified *S. cerevisiae*-specific reads. Only a small number of genes (< 10) show signs of higher expression in the multiplexed sample, indicating misallocation of reads. This could not have been a source of bias in our analysis given the high-concordance between replicates of the data, where the first replicate was not multiplexed, whereas the second was (Supplemental Fig. S5; Supplemental Table S2).

